# Development of a mobile, high-throughput, and low-cost image-based plant growth phenotyping system

**DOI:** 10.1101/2023.07.18.549560

**Authors:** Li’ang Yu, Hayley Sussman, Olga Khmelnitsky, Maryam Rahmati Ishka, Aparna Srinivasan, Andrew D.L. Nelson, Magdalena M. Julkowska

**Affiliations:** The Boyce Thompson Institute, Ithaca, NY 14850, USA

## Abstract

Nondestructive plant phenotyping is fundamental for unraveling molecular processes underlying plant development and response to the environment. While the emergence of high-through phenotyping facilities can further our understanding of plant development and stress responses, their high costs significantly hinder scientific progress. To democratize high-throughput plant phenotyping, we developed sets of low-cost image- and weight-based devices to monitor plant growth and evapotranspiration. We paired these devices with a suite of computational pipelines for integrated and straightforward data analysis. We validated the suitability of our system for large screens by evaluating a cowpea diversity panel for responses to drought stress. The observed natural variation was subsequently used for Genome-Wide Association Study, where we identified nine genetic loci that putatively contribute to cowpea drought resilience during early vegetative development. We validated the homologs of the identified candidate genes in Arabidopsis using available mutant lines. These results demonstrate the varied applicability of this low-cost phenotyping system. In the future, we foresee these setups facilitating identification of genetic components of growth, plant architecture, and stress tolerance across a wide variety of species.

## Introduction

Plant phenotyping provides a critical layer of information that helps to decipher biological processes and genetic mechanisms related to plant growth and development in response to various environmental factors (Fahlgren *et al*., 2015b; Tardieu *et al*., 2017; Tripodi *et al*., 2018; Zhao *et al*., 2019). Plant phenotypes can differ in spatial and temporal resolution, and reflect plant biochemistry, physiology, morphology, as well as agronomic performance. Plant phenotyping aids fundamental biology research and plant breeders alike through identification and development of new traits related to disease resistance, plant performance, and environmental resilience. Traditionally, plant phenotypes were collected using manual and destructive methods, associated with high experimental cost, limited throughput, and inconsistencies in data based on subjective interpretation of the observations (Furbank & Tester, 2011; Walter *et al*., 2015). Over the last two decades, image-based technologies and integration of robotics resulted in more widespread adaptation of diverse approaches to non-destructively capture plant growth, architecture and physiology (Fiorani & Schurr, 2013; Yang *et al*., 2013, 2020). These nondestructive methods have propelled plant science research forward by allowing for higher number of repeated measurements, standardization of measurements, as well as increased spatial and temporal resolution. The increased throughput of an experiment, in turn, allows for screening large populations of plants which can be further utilized in forward genetic screens (Chawade *et al*., 2019), or study the effect of various biostimulants (Rouphael & Colla, 2020). However, many phenotyping solutions still require significant monetary investment, or extensive engineering experience, which is not widely accessible at the lab, department, or even university/institute level.

The surge in availability of low-cost computers and microcontrollers, such as Raspberry Pi and Arduino, has resulted in the increased development of cost-effective phenotyping platforms, leading to greater flexibility and affordability of plant phenotyping (Jolles, 2021; Ellison Mathe *et al*., 2022; Kondaveeti *et al*., 2022). Some of the previously developed systems can be utilized to capture seed germination, shoot, and root size (Czedik-Eysenberg *et al*., 2018; Colmer *et al*., 2020; Feldman *et al*., 2021; Ohlsson *et al*., 2021; Li *et al*., 2023). While these low-cost solutions diversify image-based phenotyping, they require extensive engineering experience and equipment, and are suitable only for small plants, such as Arabidopsis, to effectively capture plant growth or provide an estimate of plant biomass. Thus, there is a growing need to develop low-cost phenotyping solutions for larger plants with more complex 3D plant architecture.

The high-throughput nature of image-based phenotyping is driven by image-processing software (Berry *et al*., 2018; Das Choudhury *et al*., 2019; Jiang & Li, 2020). The development of plant computer vision (PlantCV) was an important milestone for plant phenotyping, offering a high level of adaptability of image processing pipelines using Python scripts (Fahlgren *et al*., 2015a). The open source nature of this software and the high volume of users provides for future development opportunities, community contributions, and sustainability of the software. While PlantCV offers pipelines to analyze images being produced by RGB (Red-Green-Blue), hyperspectral, and chlorophyll fluorescence cameras (Fahlgren *et al*., 2015a), the RGB cameras remain the most widely accessible and thus have the highest application for plant research. Based on RGB images, traits such as area, convex hull, width, and height of the plant tissue can be extracted automatically, once the pixels belonging to the plant can be isolated from the background (Gehan *et al*., 2017). Phenotypes related to tissue color and disease symptoms can be extracted after parametrization of the pipeline or training with machine learning modules (Abbasi & Fahlgren, 2016). Plant architecture traits, such as number of branches, can also be extracted, however, the precision of the automated trait extraction is highly dependent on plant species and complexity (Godin, 2000). PlantCV has successfully been adopted for evaluation of a growing number of crops, including maize, rice, cassava, and more (Hairmansis *et al*., 2014; Kolhar & Jagtap, 2023). While the available image processing tools form a robust basis of image-based phenotyping (Ismael *et al*., 2022; Rossi *et al*., 2022; Zhang *et al*., 2022), most of them are characterized by strong reliance on computational user expertise. Computational pipelines often require either generating customized scripts for large-scale data processing or producing self-trained parameters for machine learning purposes. Additionally, adapting imaging hardware and software from diverse sources requires extensive optimization by end users. The programming and engineering requirement for plant phenotyping forms a substantial hindrance in the widespread application of low-cost phenotyping solutions. Thus, a low-cost phenotyping solution with integrated hardware and software, as well as the ability to accommodate a wide range of plants, is sorely needed.

In this vein, we developed an open-source system consisting of three hardware setups (**PhenoRig, PhenoCage, and AWWESmo**) and easy-to-use computational pipelines that streamline image collection and data analysis. Our system, built using low-cost and lightweight materials, can be used to monitor physiological responses in response to stress across a wide variety of species using automated RGB cameras and scales. This integrated system substantially reduces the cost and time necessary to collect reproducible image and evapotranspiration data, and lowers the computational barrier to extract phenotypic data (**RaspiPheno Pipeline**). Moreover, the developed ShinyApp (**RaspiPheno App**) is a dynamic tool for downstream statistical data analysis, sample comparison, and data interpretation. To showcase the system’s capacity, we used the developed tools to screen a natural diversity panel of 368 cowpea genotypes and, through Genome-Wide Association Study (GWAS), identified new genes associated with drought response during early vegetative growth. The described suite of phenotyping solutions, as well as data analysis pipelines, will promote affordable plant phenotyping and accelerate the discovery of new genes and physiological traits contributing to stress resilience.

## Materials & Methods

### Phenotyping facility development

#### Top-view imaging of Arabidopsis using PhenoRig

To continuously collect top-view images, we built a PhenoRig where 32 pots (width: 2.5 inch ×length: 2.5 inch×height: 3.5 inch for each insert), each containing a single Arabidopsis plant, could be imaged simultaneously within one setup. We built a wooden frame (length: 60 × width : 40 × height 43.5 cm for the frame) to accommodate a standard 1020 (∼27.94 x 54.61 cm) rectangular tray. The images were collected by two Raspberry Pi cameras connected to a RaspberryPi Zero computer through an Arducam Multi-camera adapter, with flex cables. To prevent the movement of plants during plant watering, a plastic tray was anchored to the bottom of the PhenoRig. The imaging of plants was performed every 30 minutes using the automated startup script (described in pheno-computational pipeline development section below). No additional light source was installed, and the images collected during the night period were removed in the automated data preparation workflow by specifying the start and end time of the light period (hour-minute format: HH.MM) of image collection each day as a parameter input for data allocation (https://github.com/Leon-Yu0320/BTI-Plant-phenotyping). The materials necessary to build a PhenoRig are listed in **Supplemental Table 1**.

#### Side-view imaging using PhenoCage

To adequately capture the digital biomass of the plants with complex 3-D architectures, such as bean plants, we developed a side-view imaging PhenoCage platform. Within a cage (length: 90 cm × width: 60 cm × height: 60 cm), we placed a rotating platform on which a plant pot is placed. The background noise is limited by white semi-light-permeable plastic sheets attached to the frame. The plant is illuminated from four sides and the top using LED light bars to eliminate shading. To ensure that all plants will be positioned in the same way on the rotating platform, we attached the pot that has been cut in half in the horizontal direction to the rotating platform. The plants are imaged using one Raspberry Pi camera connected to a Raspberry Pi 4. A household shell script (PhenoCage_capture.sh) takes seven consecutive images, representing a side view of the imaged plant taken every 51.4° (https://github.com/Leon-Yu0320/BTI-Plant-phenotyping/tree/main/data_acquisition). The materials necessary to build a PhenoCage are listed in **Supplemental Table 2**. Accessories to hold the RaspberryPi and monitor were designed using TinkerCad (https://www.tinkercad.com) and produced by a 3-D printer (PRUSA i3 MK3S+). Examples and details of the 3-D printed components can be found at https://github.com/Leon-Yu0320/BTI-Plant-phenotyping/tree/main/3Dprint.

#### Monitoring evapotranspiration using AWWEsmo

To monitor the plant evapotranspiration, we developed an Arduino-Watering and Weighing unit to measure Evapotranspiration (AWWEsmo). Using this device, the pots are placed on a scale, where they are automatically weighed and watered to their target weight. The Arduino controller was connected to a tensiometer that served as a scale, and to a submerged 5-volt pump that was activated for a period calculated to be necessary to reach the target weight. The controller and the mini-breadboard were placed in a 3D-printed container, designed to protect the electrical components from splatter and dust in the growth chamber. The tensiometer constituting the scale was attached to the saucer, to prevent water spillage onto the electrical components, and housed in a 3D-printed design, allowing leveling of the scale and support for the hose connected to the pump. The materials necessary to build an AWWEsmo device are listed in **Supplemental Table 3**, while the detailed instructions on how to build and program the device can be found at https://github.com/ok84-star/AAWSMO. Details on the 3D-printed designs can be found at https://github.com/Leon-Yu0320/BTI-Plant-phenotyping/tree/main/3Dprint/AWWESmo. The detailed usage manual, including calibration and execution of the AWWEsmo, are available at protocols.io(Julkowska *et al*., 2019; Khmelnitsky *et al*., 2023). Supplemental Video on assembly can be found here: https://youtu.be/QlJUVdQT6VA.

#### Pheno-computational pipeline development

The plant image collection is integrated into a customizable shell script, optimized for ISO, image sharpness (sh), contrast (co), brightness (br), shutter speed (ss), size of the image for individual imaging conditions. For PhenoRig setup, the automated data collection was conducted with specifications of time interval (**unit: min**), duration (**unit: days**), and hardware identifiers (**format: RaspiID_cameraID**). Once the image and experimental settings are determined by the user, the imaging command is deployed at determined time intervals using crontab which is nested within the setup scripts. Users can launch the collection by using a local Raspi computer, or connect Raspi computers to the internet and launch the program remotely by a personal computer (PC) or a server. For PhenoCage setup, image data collection is launched manually for individual plants. For each plant, images from seven sides were collected with a hardware identifier (**format: RaspiID**) and side numbers (**format: side1-side7**). After each experiment session for both PhenoRig and PhenoCage, images can be transferred using USB flash drive or using ssh transfer proxy to a server or other local devices.

After image collection, the pipeline requires experiment-specific parameters as input to execute the image analysis correctly. The parameters guide key steps related to image transformation, masking, selection of regions of interest, and extraction of phenotypic data into an image analysis protocol derived from plantCV image processing algorithm (Gehan *et al*., 2017), a tutorial for parameter setup for plantCV software was instructed using example images collected with PhenoRig and PhenoCage (Yu & Julkowska, 2022). The computation pipelines and RasPiPheno pipeline are publicly available on Github with manual and supplementary information provided (https://github.com/Leon-Yu0320/BTI-Plant-phenotyping).

The quantitative data obtained from collected images are subsequently analyzed for changes in digital biomass throughout time/treatment. Prior to statistical data analysis, data collected using PhenoCage is additionally processed by summarizing the pixels representing shoot projected area from 7 side view images. Once the digital biomass of each plant is determined from either side-view or top-view images, the genotype, replicate, and treatment information is decoded using a meta-data table (https://github.com/Leon-Yu0320/BTI-Plant-phenotyping/tree/main/Results_example).While PhenoRig images are decoded based on positional information, the PhenoCage data is decoded based on the timestamp of the image, assuming that the experiment is imaged sequentially in order of the pot position. The decoded data is subsequently processed under the framework **RasPiPhenoApp** (https://github.com/Leon-Yu0320/BTI-Plant-phenotyping), a web interactive and streamlined analysis tool. Using the smooth spline, loess fit, or polynomial regression fit functions, each data point is curated to generate curated values as smoothed dataset. The original datapoints that exceed the 1∼3 times standard deviation (SD) relative to corresponding curated values can be classified as outliers. The user can remove specific points based on the customized cutoff (1 to 3 times SD), to generate a clean dataset. The growth rates (GR) are calculated using a linear function either for the entire duration of the experiment (PhenoCage) or for each day of the experiment (PhenoRig). The differences between treatments and/or genotypes (or in other single factors, two factors experiments) are subsequently tested using t-test, Wilcox, ANOVA, or two-way ANOVA regarding the curated plant leaf area, the leaf area without outliers, and growth rate The visualization of the graphs is performed using ggplot2 and ggpubr packages (Gentleman *et al*., 2016). The integration of data analysis tools into a graphical user interface is performed using *shiny* R package (https://shiny.rstudio.com).

### Plant Material and Plant Growth Conditions

#### Arabidopsis thaliana

Arabidopsis (*Arabidopsis thaliana*) Col-0 seeds were sterilized for 10 min with 50% bleach and rinsed five times using Milli-Q water and germinated on ½ strength x Murashige and Skoog (MS) medium containing 0.5% (w/v) sucrose, 0.1% (w/v) 4-morpholineethanesulfonic acid (MES), and 1% (w/v) agar. After 24 h of vernalization at 4℃ in the dark, the plates were placed in the Conviron growth chamber with the light intensity of 130–150 µmol m^−2^ s ^−1^ in a 16 h light / 8 h dark cycle at 21℃ and 60% humidity. At 7 days after germination, the seedlings were transplanted to soil (Cornell Mix) watered to 100% soil water-holding capacity and placed in a walk-in growth chamber with the abovementioned conditions. When the pots dried to the weight corresponding to 50% of their water holding capacity, they were soaked for 1 h in tap water or a 200 mM NaCl solution, resulting in a concentration of 100 mM NaCl based on the 50% soil water holding capacity, which corresponded to a moderate level of salt stress according to (Awlia *et al*., 2021). We allowed the pots to be drained for 2-3 h to eliminate excess moisture. The pots were subsequently placed under PhenoRigs equipped with an automated imaging system, and the pot weight was measured and adjusted daily to maintain the reference weight corresponding to 50% of the soil water holding capacity throughout the experiment. At the end of the experiment, fresh weight was collected for all imaged plants. The collected images were processed using the pheno-computational pipeline described above, and the data were processed in R **(Supplemental Table S4).**

#### Cowpea pilot experiment

The seeds of 5 cowpea accessions (CB5-2, IT97K-499, Sanzi, Suvita-2, UCR799), representative of the wider diversity within cowpea were germinated in 4-inch pots filled with soil (Cornell Mix + Osmocote)(Liang *et al*., 2023). We used seven biological replicates per accession per treatment for this experiment. The control and drought-treated plants were kept at 60% and 20% soil water-holding capacity, respectively. To determine target weights for each pot, we left the pots to air dry for 72 hours, and assigned this weight to correspond to 0% water-holding capacity. We then saturated the soil for 24 hours, removed excess water, weighed the pots, and assigned this value as the 100% soil water-holding capacity weight. At this point, we sowed two seeds per pot, and thinned to one seedling per pot after germination occurred. We initiated tracking pot weight at seventeen days after germination, watering each pot to its target weight daily for fifteen consecutive days. Drought treatment target weights were reached on day four after tracking started. We imaged the plants using the PhenoCage setup starting at seventeen days after germination and subsequently every other day for the next two weeks (resulting in total of seven time-points, with each time-point consisting of 7 images for each plant). At the end of the experiment, the fresh weight of the cowpea shoot was collected for all the imaged plants. The collected images were processed using the pheno-computational pipeline described above, and the data was processed in R **(Supplemental Table S4).**

#### Tepary bean pilot experiment

The seeds of 2 tepary bean accessions (TDP359 and TDP22), representative of the wild and cultivated tepary bean respectively (Muñoz-Amatriaín et al., 2021), were germinated in 4-inch pots filled with soil (Cornell Mix + Osmocote) watered to 100% soil water-holding capacity. We used 6 biological replicates of TDP359 (cultivated) and 12 replicates of TDP22 (wild) per treatment for this experiment. The control and drought-treated plants were kept at 60% and 10% soil water-holding capacity, respectively. The drought was imposed as described for the cowpea pilot experiment above. We imaged the plants using the PhenoCage setup starting at seventeen days after germination and repeated every second day for the consecutive two weeks. At the end of the experiment, the fresh weight of the tepary bean shoot was collected for all the imaged plants. The collected images were processed using pheno-computational pipeline described above, and the data was processed in R **(Supplemental Table 4).**

### Cowpea mini-core population screen

The cowpea mini-core population, consisting of 368 accessions (Muñoz-Amatriaín *et al*., 2021) was screened as described for the cowpea pilot experiments. The accessions were distributed over six experiments, and we used five accessions (CB5-2, IT97K-499, Sanzi, Suvita-2, UCR799) as internal standards for each experiment. One accession, TVu-9393 was excluded because it did not germinate after multiple trials, and another accession, TVu-3965, was omitted due to lack of seeds available. The pot weight was monitored and adjusted every day, while imaging of the plants using PhenoCage was performed every 2^nd^ day. Due to the various growth habits of cowpea, we occasionally added transparent, 3D-printed support to ensure the upright position of the plant. The weight of the support was accounted for in the evapotranspiration data analysis. Additionally, we measured photosynthetic efficiency, leaf temperature and chlorophyll content using PhotoSynQ device 6 and 13 days after treatment initiation. At the end of each experiment, the fresh weight of the cowpea shoot tissue was collected for all the imaged plants. The collected images were processed using the pheno-computational pipeline described above. This dataset includes six sets of experiments, evaporation rate curation for individual plants, and the side-view image data comparison derived per experiment was performed using the R scripts (**Supplemental Table 4)**. Subsequently, individual experimental data were merged, modeled using smooth splines, used to calculate growth rate and cumulative evapotranspiration per gram of fresh weight, and prepared for subsequent GWAS (**Supplemental Table 4)**. The raw and curated data can be accessed in open-access Zenodo Repositories (overview and links are listed in **Supplemental Tables 5 - 7**).

### Genome-wide Association Study of drought stress responses in cowpea

All collected and curated phenotypic data was used for GWAS. The kinship matrix was calculated for all included accessions using GAPIT (Wang & Zhang, 2021), and included as a co-factor in the GWAS mixed model (https://github.com/arthurkorte/GWAS). The GWAS model uses fast approximation (Zhang & Liu, 2011) and relies on the ASReml library (Butler *et al*., 2009). The results files were subsequently processed to draw QQ-plots, indicating any bias within the GWAS model, Manhattan plots to identify significant associations above the Bonferroni threshold, as well as the effect size plots to evaluate the estimated effect size of the loci selected for further inspection (**Supplemental Table 4**). The identified genomic regions were compared between the traits mapped under control and drought stress conditions (**Supplemental Table 8**). Loci identified exclusively under drought stress conditions were considered for further evaluation. The GWAS output files, as well as all the generated plots, can be accessed in open-access Zenodo Repository (https://doi.org/10.5281/zenodo.7438567).

### Screening of homologs in Arabidopsis for their contribution to drought resilience

The drought-specific loci identified through cowpea GWAS described above were investigated for annotated genes within the linkage disequilibrium (LD, 30 kbp) of the identified SNP. Arabidopsis sequence homologs to the genes within the LD were acquired from the cowpea genome annotation (Lonardi *et al*., 2019). For each identified Arabidopsis homolog, we explored publically available homozygous T-DNA insertion lines that exclusively target our gene of interest. The lines and their corresponding cowpea genes are listed in **Supplemental Table 9**. The seeds were ordered from ABRC (https://abrc.osu.edu/), and the seeds of each mutant line were grown alongside Col-0 genotype, as described for the Arabidopsis phenotyping experiment above. Two weeks after germination, the seedlings were exposed to control or drought stress conditions (60 and 10% of soil water-holding capacity, respectively). The plants were monitored for growth using the PhenoRig setup every 30 minutes, while evapotranspiration of every plant was monitored every 48 hours using the AWWEsmo device. Based on the results and phenotypes of mutants with significantly affected growth rate under drought stress, we made a selection of 14 T-DNA insertion lines for further evaluation (CP.GR4-1, CP.GR4-2, CP.NPQ6-1, CP.NPQ6-2, CP.NPQ6-3, CP.NPQ6-4, CP.NPQ6-5, CP.EVT2-1, CP.EVT2- 2, CP.EVT3-1, CP.EVT3-2, CP.EVT6-1, CP.EVT6-2, CP.EVT8, **Supplemental Table 9**). 100% water holding capacity was determined as described above for the Cowpea Pilot Experiment. Concurrently, we grew the 14 T-DNA insertion lines on ½ MS plates for 10 days and transferred the seedlings to soil. Two weeks after germination, we initiated tray imaging every 30 minutes. To efficiently bring the drought treatment pots down to 20% water holding capacity, we placed small fans above them for a 90-minute increment. Seventeen days after germination, we began tracking water use every second day, and this day was marked as day 1 of stress. Images were taken for two weeks. Primary bolts were cut from plants that began flowering within these two weeks to prevent bias in the image analysis. The data was analyzed in the same way as described in previous experiments. The specific R markdown files and raw dataset can be accessed on Github page (https://github.com/Leon-Yu0320/BTI-Plant-phenotyping/tree/main/ImageData_curation_example).

## Results

### Phenotypic hardware design of the system

To increase the accessibility of plant phenotyping, we developed a set of mobile, affordable, and customizable phenotyping setups. The setups were designed to fit into conventional growth chambers (**Fig. 1**) and allow the evaluation of plants with different types of architecture. The top-view **PhenoRig** setup was designed to image the growth of plants for which the majority of growth occurs within two dimensions, such as Arabidopsis rosettes (**Fig. 1A**). Plants with more complex architecture can be imaged using side-view imaging, as possibilities for top-view imaging are often limited by the vertical space available within a typical growth room. In the side-view **PhenoCage** setup (**Fig. 1B**), a plant with a more complex 3D architecture is positioned on a rotating platform, and seven consecutive images are taken to adequately capture the projected shoot area and reflect the plant’s digital biomass. To monitor plant evapotranspiration, we developed an Automatic Weighing and Watering device to study Evapotranspiration (**AWWEsmo**) that automatically records pot weight and waters it to the reference weight (**Fig. 1C**).

**Figure 1.**
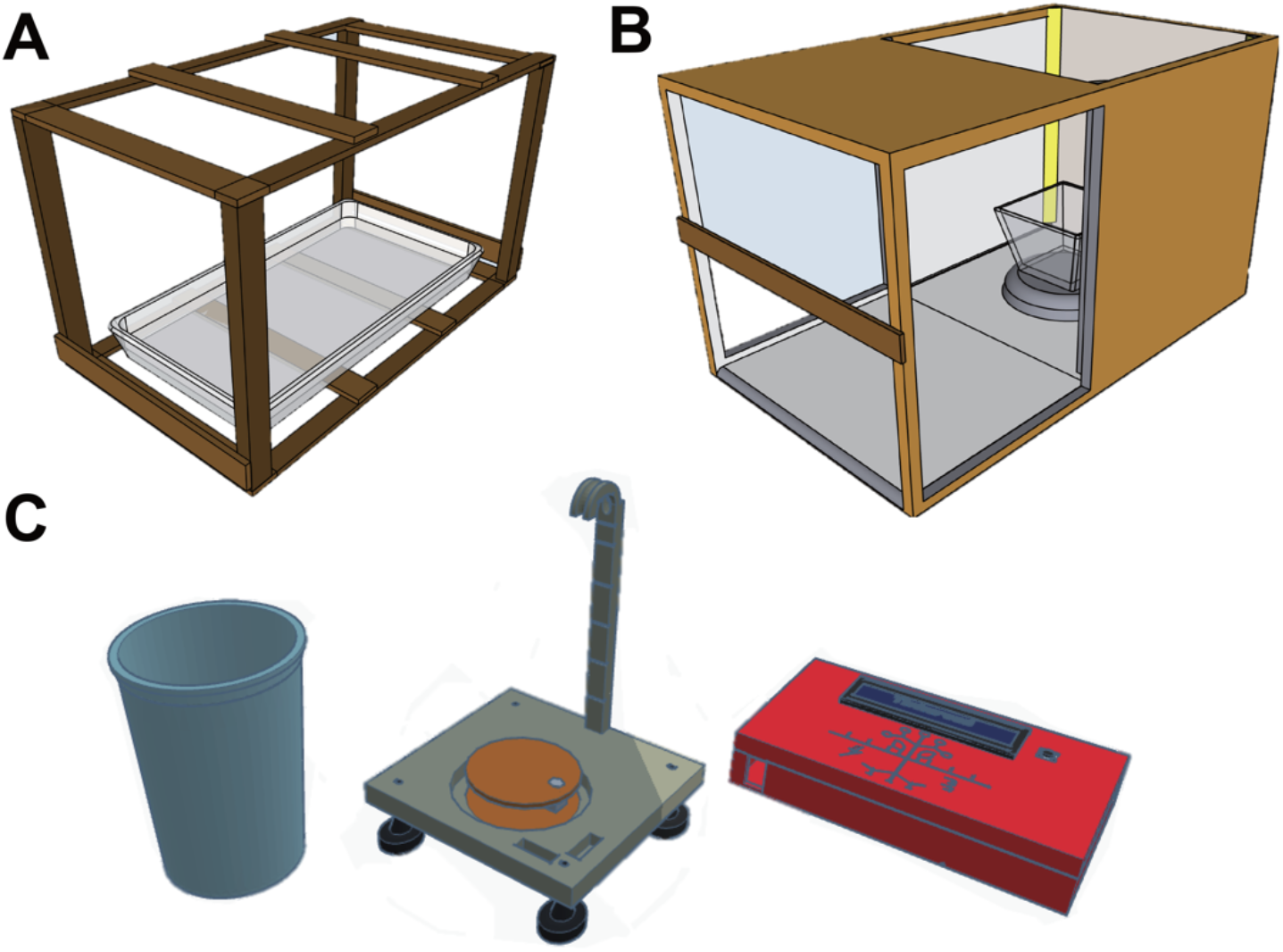
PhenoRig, PhenoCage, and AWWEsmo Design: Three facilities were constructed using lightweight materials with 3-D printed accessories for data collection purpose. **(A)** PhenoRig **system:** including a frame to hold two Raspi cameras and a Raspi computer. A tray holding plants is placed at the bottom of the frame to display plants’ top-view characteristics. **(B) PhenoCage system:** including a frame to hold one Raspi camera and a Raspi computer. A rotating booth is placed at the center of the cage with a constant rotation rate to represent plant architectures from different sides. **(C)** AWWESmo system: including a container for water replenishment (blue), and a water pump that integrates a weight scale controlled by an Arduino Uno R3-derived console (red). Precise water replenishment is executed by the input weight (grams) of the console.

The individual devices rely on Raspberry Pi computers (**PhenoRig and PhenoCage**) and Arduino microcontrollers (**AWWEsmo)** for data acquisition, which lend themselves to flexible and cost-effective setups that can be easily adapted to accommodate a wider range of species. The PhenoCage can be used to monitor the growth of Arabidopsis continuously, using an automatically deployed imaging command, while PhenoRig and AWWEsmo currently require the user to feed the plants into the setup and deploy the image/measure command manually. The current design of PhenoRig allows imaging of a standard full tray of Arabidopsis plants with two cameras, where each camera can capture a grid of 4×4 plants (**Fig. 2A**), with the total capacity of PhenoRig being an 8×4 plant grid. PhenoCage, on the other hand, has a capacity of one plant, as the complex 3D architecture of the shoot does not permit simultaneous imaging of multiple plants. To ensure the best results in image processing, we recommend putting 2-4 white tags on top of the pots for PhenoRig, to correct for white balance between the individual images (**Fig. 2B**). For the PhenoCage, we suggest using a white background for white balance corrections (**Fig. 2A, 2B**). Additional accessories installed for image collection and illumination, include RasPi cameras LED lights, LCD touch screens, and 3D-printed accessories holders (**Fig. S1, Fig. S2)**

**Figure 2.**
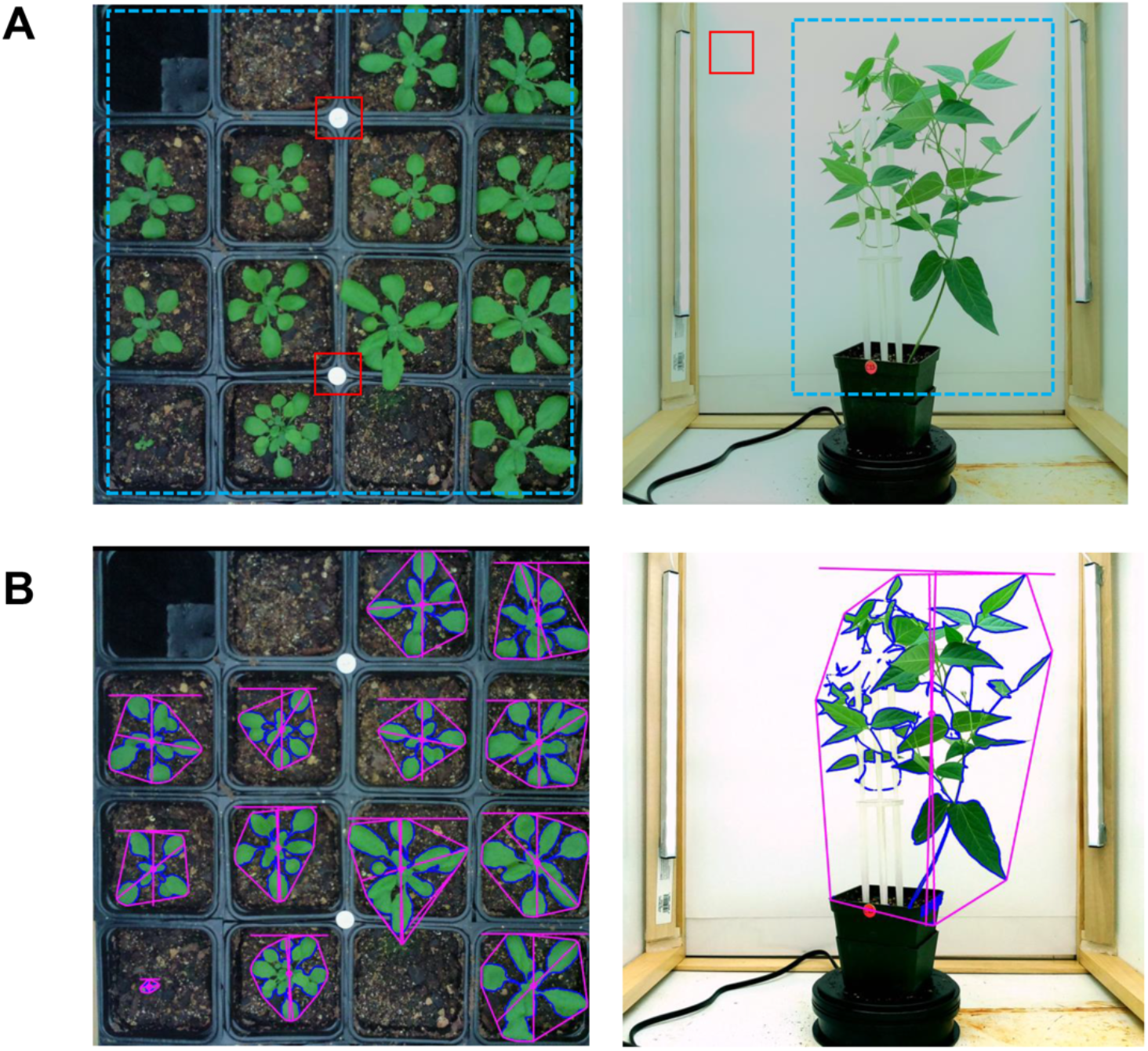
PhenoRig and PhenoCage images. **(A)** RGB images output of Raspi camera of Arabidopsis (left panel) and tepary bean (right panel). The red squares highlight the area used for white balance correction in the subsequent steps. Blue squares indicate the region of interests (ROIs) for image processing. **(B)** Output images after processing using PlantCV pipeline. The blue edge highlights the perimeter of the leaf area (green tissue) and the purple edge defines the convex hull area of each individual plants.

To effectively address data analysis of images collected over multiple days and devices, we developed a computational pipeline (**RaspiPheno Pipeline**) that automates image acquisition, storage, and subsequent data extraction into a repeatable workflow (**Fig. 3**). This pipeline parallelizes image segmentation steps and deploys them on batches of images to examine projected shoot area and architectural traits (**Fig. 3**). This optimized image extraction process requires 1) the positional information of white balance markers and region of interest (ROI, **Figure 2**), 2) specific cut-offs and coordinates used for extracting plant objects from RGB images, and 3) storage locations for input and output files (**Fig. 2B**, **Fig. 3**). Quantitative data generated and organized by the RasPiPheno pipeline can be analyzed using **RaspiPheno** App, an interactive and programming-free analytical application powered by Shiny R. RaspiPheno App was designed to address statistical analysis of data associated with shoot area and architectural traits among customized independent variable groups (eg: genotypes or treatments). Within RaspiPheno App the user can match the information on genotype and treatment with quantitative values from each individual plant as the data reshaping process (**Fig. 3**). The integrated data are then presented as time-series graphs, and the user is presented with data curation function to smooth noisy data and generate a predicted, or transformed dataset (see **Methods**). Alternatively, the user can generate a clean dataset by removing datapoints that are beyond the standard deviation (SDs) of values predicted by the smoothing function. RaspiPheno App can calculate growth rate for the user within a customized time interval (eg: 12 or 24 hours) to characterize the differences of plant growth under different conditions using pairwise or multiple-group tests **(Figure 3**). More information on using RaspiPheno can be found at (https://github.com/Leon-Yu0320/BTI-Plant-phenotyping/tree/main/RasPiPheno_APP).

**Figure 3.**
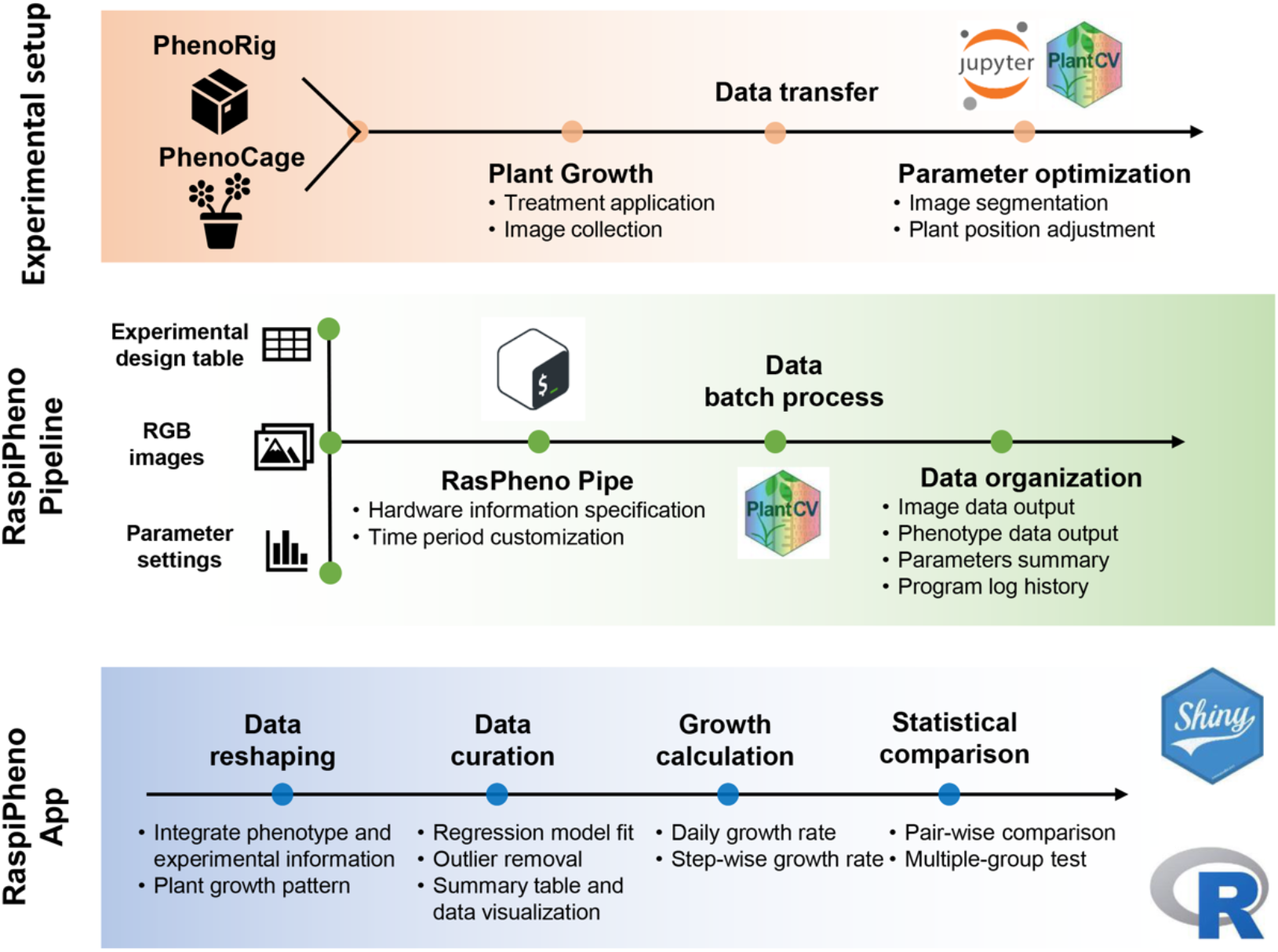
Overview of the Pheno-Computational pipeline. The experimental setup consisting of PhenoRig (Arabidopsis) or PhenoCage (larger plants with 3D architecture is used to collect data on plant growth and architecture. Using the first few images obtained by the experimental setup system, the image acquisition parameters are optimized for best image processing in the subsequent steps. The phenotypic data processing is based on PlantCV pipeline integrated into **RaspiPheno Pipeline**, which applies batch-specific parameters (*parameter settings table*) onto a batch of *RGB images* (indicated in *experimental design table*) to perform batch-specific processing. Subsequently, the quantitative data from RasPiPheno pipeline can be uploaded along with the metadata, containing genotype and treatment information for individual plants, into **RasPiPheno App**, where the data is matched, curated, processed for outliers and custom calculations of plant growth rate and statistical comparisons are performed.

Together, the RaspiPheno pipeline and RaspiPheno App provide an integrated framework for extracting data from images and quickly analyzing the phenotypic data. As a web-browser integrated RShiny application, the RaspiPheno App streamlines what is typically a command line-based statistical analysis into an intuitive and interactive process. As the developed tools have limited computational requirements, they can be run on a standard laptop with an internet connection. As a result, we envision these open-source hardware and software packages simplifying the data extraction process that often serves as a roadblock in any bioinformatic analysis.

The instructions for constructing system hardware using inexpensive wooden or aluminum frames can be accessed at (Yu & Julkowska, 2022), whereas the necessary parts for constructing each setup are listed in (Supplemental Tables 1, 2, and 3). The RaspiPheno App and RaspiPheno Pipeline are available, along with the detailed instructive user manuals and example datasets, on GitHub repository (https://github.com/Leon-Yu0320/BTI-Plant-phenotyping). Together, PhenoRig, PhenoCage, and AWWEsmo represent a basic suite of plant phenotyping tools that significantly accelerate research and can be instrumental in screening populations of accessions or mutants under diverse conditions. The construction of each setup requires minimal financial investment (less than U.S. $200), and thus further contributes to democratizing plant phenotyping tools in a wide range of potential users.

### Stress-induced changes in Arabidopsis, cowpea, and tepary beans

To evaluate the efficacy of the developed tools, we evaluated if images derived from the system are of sufficient quality to detect the effect of abiotic stress on three species - Arabidopsis, cowpea, and tepary beans (**Fig. 4**, **Fig. 5**). Arabidopsis Col-0 plants were treated with salt stress (100 mM NaCl effective concentration) at two weeks after germination and imaged every 30 minutes for the following two weeks. As expected, we observed a consistent decrease in rosette size starting from 6 days after induction of salt stress (**Fig. 4A**).

**Figure 4.**
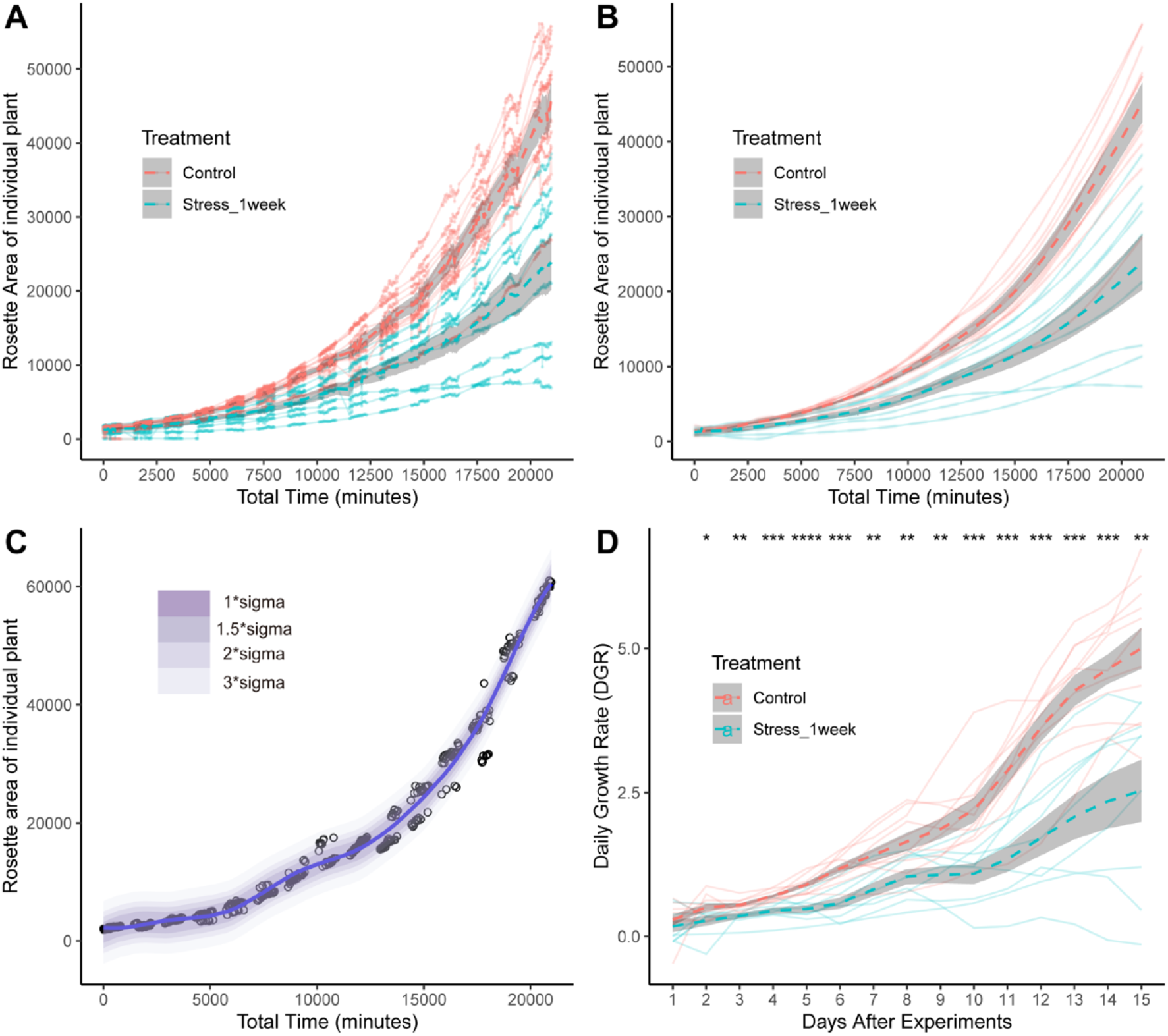
Example of Arabidopsis PhenoRig data. Arabidopsis seedlings were exposed to salt stress two weeks after germination. The trays containing Col-0 seedlings exposed to salt stress / mock treatment (Control) were placed under PhenoRig setup and imaged every 30 minutes for 2 weeks. **(A)** The increase in projected rosette area was observed over 2 weeks following the salt stress exposure. **(B)** In order to reduce the noise in data, we curated data for each individual plant using smooth spline function with different level of sigma to identify data points as potential outliers. **(C)** Data derived from smooth spline function was calculated for all the samples used for imaging within our experiment, which significantly reduced noise levels. **(D)** The smooth spline data was subsequently used for calculating daily growth rate for each measured plant. Difference in daily growth rate between plants exposed to control and salt stress treatment was calculated using t-test. The *, **, *** and **** represent p-value below 0.05, 0.01, 0.001 and 0.0001 respectively.

**Figure 5.**
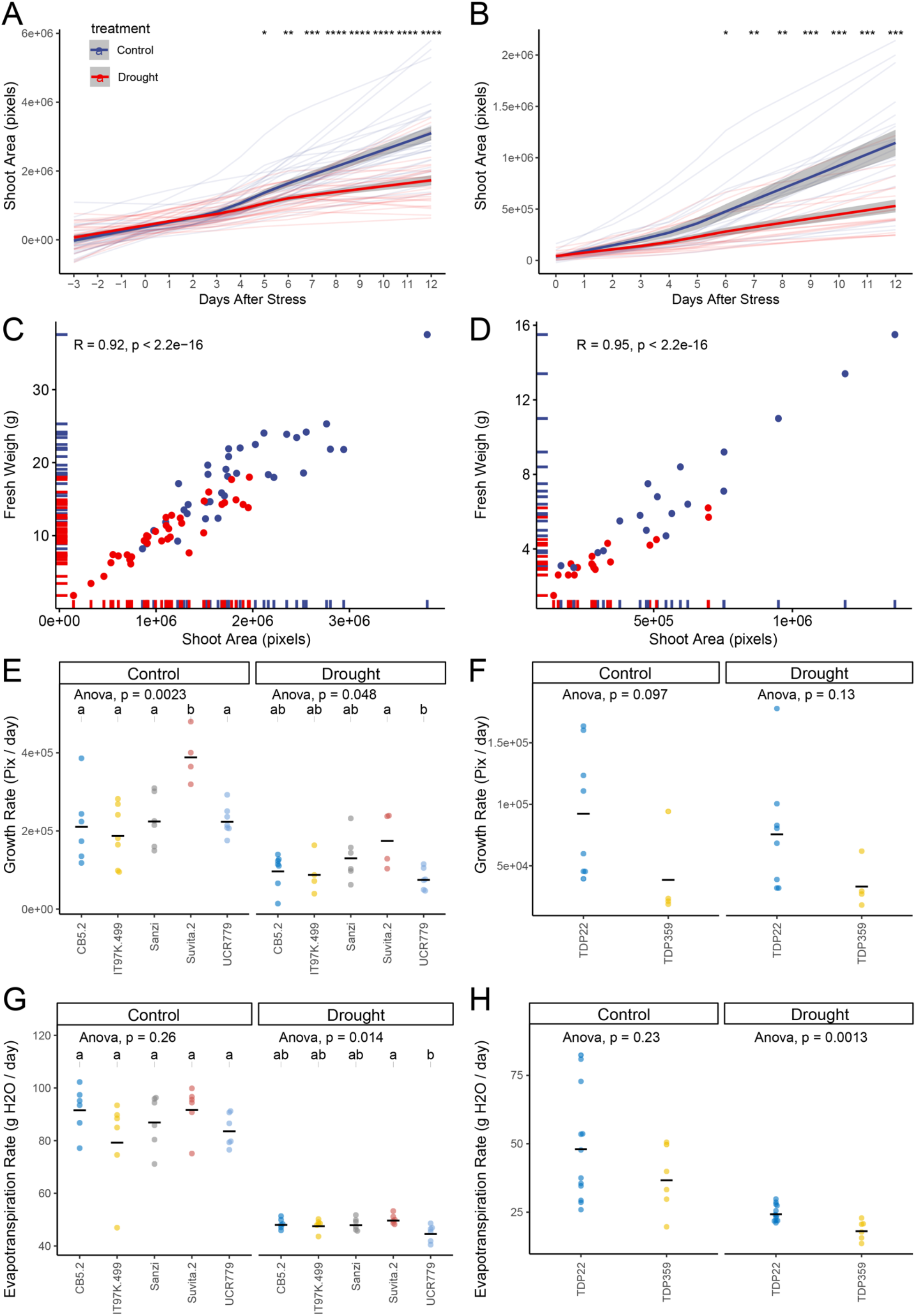
Examples of cowpea and tepary bean data collected using PhenoCage and AAWEsmo setups. The seedlings of 5 cowpea accessions, and 2 tepary bean accessions were germinated for 17 days, and exposed to control treatment or drought stress (60% and 10% soil water-holding capacity respectively). **(A)** The increase in shoot area was modeled using the smooth spline function over the recorded data for cowpea and **(B)** tepary bean over the course of 12 days following the drought stress application. The difference between treatments was calculated using ANOVA with *, **, *** and **** indicating the p-values below 0.05, 0.01, 0.001 and 0.0001 respectively. The Pearson correlation between the projected shoot area and fresh weight of the shoot recorded at the last day of the experiment was examined in both **(C)** cowpea and **(D)** tepary bean. The growth rate was calculated based on the smooth spline modelled data for **(E)** 5 cowpea accessions and **(F)** 2 tepary bean accessions for individual conditions. The median evapotranspiration rate, was calculated based on the data collected using AAWEsmo for **(G)** 5 cowpea accessions and **(H)** 2 tepary bean accessions. The effect of the genotype within individual treatment was tested using ANOVA, and the significantly different groups of cowpea accessions were additionally determined using Tukey HSD test (p-value < 0.05).

While continuous imaging can provide highly detailed information, it is also prone to variation due to leaf movement throughout the day. To reduce this variation, the rosette size data was modeled over the entire experimental time course, with images taken during the dark removed, using a smooth spline function within PhenoApp (**Fig. 4B**). These splines provide the means for smoothing noisy data points by providing function estimates, balancing a measure of goodness of fit with a derivative based measure of the smoothness. This function was also used to identify potential outliers and eliminate specific points from the data set based on standard deviations from the spline (**Fig. 4C**). Plotting the increase in the rosette size of individual plants using smooth splines significantly reduced the noise caused by diurnal movements of leaves, and thus provides a clearer image of the plant’s growth trajectory. We used the collected data to calculate the daily rosette growth rate by fitting a linear regression to daily changes in rosette size and plotting the change in growth rate over the course of the experiment. The daily growth rate decreased significantly within two days of salt treatment application (**Fig. 4D**). The difference between non-stress and salt-stressed plants in growth rate increased over the duration of the experiment (**Fig. 4D**). These results suggest that our PhenoRig setup allows us to identify differences between plants grown under non-stress and salt stress conditions with high sensitivity, detecting significant differences as early as two days after stress in daily rosette growth rate.

To evaluate the efficacy of the PhenoCage setup for more complex plant architectures, five cowpea accessions and two tepary accessions were exposed to drought stress at 17 days after germination. For the two weeks after stress application, the weight of each pot was monitored and adjusted daily using AWWEsmo, while changes in shoot size were recorded every two days using the PhenoCage setup (**Fig. 5**). As some plants required additional support structure, due to their climbing or prostrate growth habit, we designed a stackable trellis that was 3D printed using a transparent filament, which resulted in minimal obstruction of the imaged plant area (**Fig. S3**). Drought stress was applied through a gradual reduction in soil water-holding capacity from 60 to 10% for both cowpea and tepary beans. The differences between control and drought stress plants were observed after five and six days, respectively (**Fig. 5A-B**). We observed high correlation between the plant fresh weight and projected shoot area recorded on the final day of the experiment for both cowpea and tepary beans (**Fig. 5C-D**), indicating high reliability of the PhenoCage system to non-destructively estimate the changes in digital biomass. When we calculated growth rate for each plant throughout the entire experiment, we observed significant differences between genotypes and treatments for tepary bean and cowpea alike (**Fig. 5E-F**). The weight of the pot and watering data, collected through AWWEsmo, was used to calculate daily evapotranspiration rate for each plant. As the target drought weight was reached 2 days after initial treatment application, the differences in evapotranspiration were also evident within 2 days after monitoring soil water holding capacity (**Fig. S4**). Evapotranspiration of tepary beans and cowpeas substantially decreased in response to drought stress (**Fig. 5G-H**). High variability in cultivated tepary bean growth rate and evapotranspiration is linked to plant size (**Fig. 5F, H**). When comparing the median evapotranspiration per plant throughout the entire experiment (**Fig. 5G-H**), significant differences were only observed under drought stress conditions. Cowpea accessions Suvita-2 and UCR779 showed the highest and lowest evapotranspiration under drought stress, respectively. Cultivated tepary bean accession (TDP-22) showed higher rate of evapotranspiration compared to the wild tepary bean accession (TDP-359). These results indicate that PhenoCage and AWWEsmo can detect not only differences between treatments but also subtle differences in plant growth rate and evapotranspiration between genotypes for plants with complex architecture, such as cowpea and tepary beans.

### Drought-stress induced changes in the cowpea diversity panel

To illustrate the suitability of the developed system for a true high-throughput phenotyping experiment, we screened a cowpea miniCore diversity panel (Muñoz-Amatriaín *et al*., 2021), consisting of 368 accessions, for drought stress-induced changes in growth rate, evapotranspiration, and photosynthetic efficiency. One replicate per accession per treatment was germinated in well-watered conditions. Once 80% of the plants developed the first trifoliate leaf, pot weight was monitored and adjusted to target weights, corresponding to 60% and 20% of soil water-holding capacity for control and drought stress, respectively. Daily evapotranspiration was monitored for fourteen days, with digital plant biomass collected every other day with the PhenoCage. Additional measurements on photosystem II efficiency were collected from each plant at 6 and 13 days after stress application. At the end of the experiment, fresh weight data were collected from shoot material for each plant.

As in pilot experiments, the high correlation between fresh weight and projected shoot area (**Fig. 6A**) indicated that our PhenoCage system produces a good estimate for digital plant biomass. Tracking progression in shoot size allowed shoot growth to be modeled using smooth splines, revealing significant differences in shoot size starting from four days after initial drought stress application (**Fig. 6B**). Based on the increase in shoot area, growth rate was also estimated for each genotype, with significant differences observed between control and drought stress conditions (**Fig. 6C, Supplemental Table 6**). Additionally, relative impact of stress on growth rate was calculated for each genotype by dividing the genotypic mean growth rate observed under drought stress conditions by the genotypic mean growth rate observed under control conditions (**Fig. 6D, Supplemental Table 6**). While on average, growth rate was reduced to 0.6 of the rate observed under control conditions, 19 accessions displayed increased vigor under drought stress (relative growth rate > 1, **Supplemental Table 6**), while 64 accessions showed severe drought stress sensitivity (defined as relative growth rate < 0.4, **Supplemental Table 6**). Drought-treated plants transpired significantly less water than control plants (**Fig. 6E**), and on average, the evapotranspiration decreased to 0.55 of the levels observed under control conditions (**Fig. S5, Supplemental Table 7**). While drought stress was observed to reduce quantum yield (Fv’/Fm’) only during the early phase of the experiment (**Fig. 6F, Supplemental Table 7**), a decrease in chlorophyll content (SPAD) was observed only at the later stage of the experiment (**Fig. 6H, Supplemental Table 7**). On the other hand, the drought stress significantly increased leaf temperature both at the early and late stage of drought stress treatment (**Fig. 6G**). These results illustrate that variability in drought stress responses across a large and diverse panel of plants with complex architecture can be captured through our PhenoCage setup.

**Figure 6.**
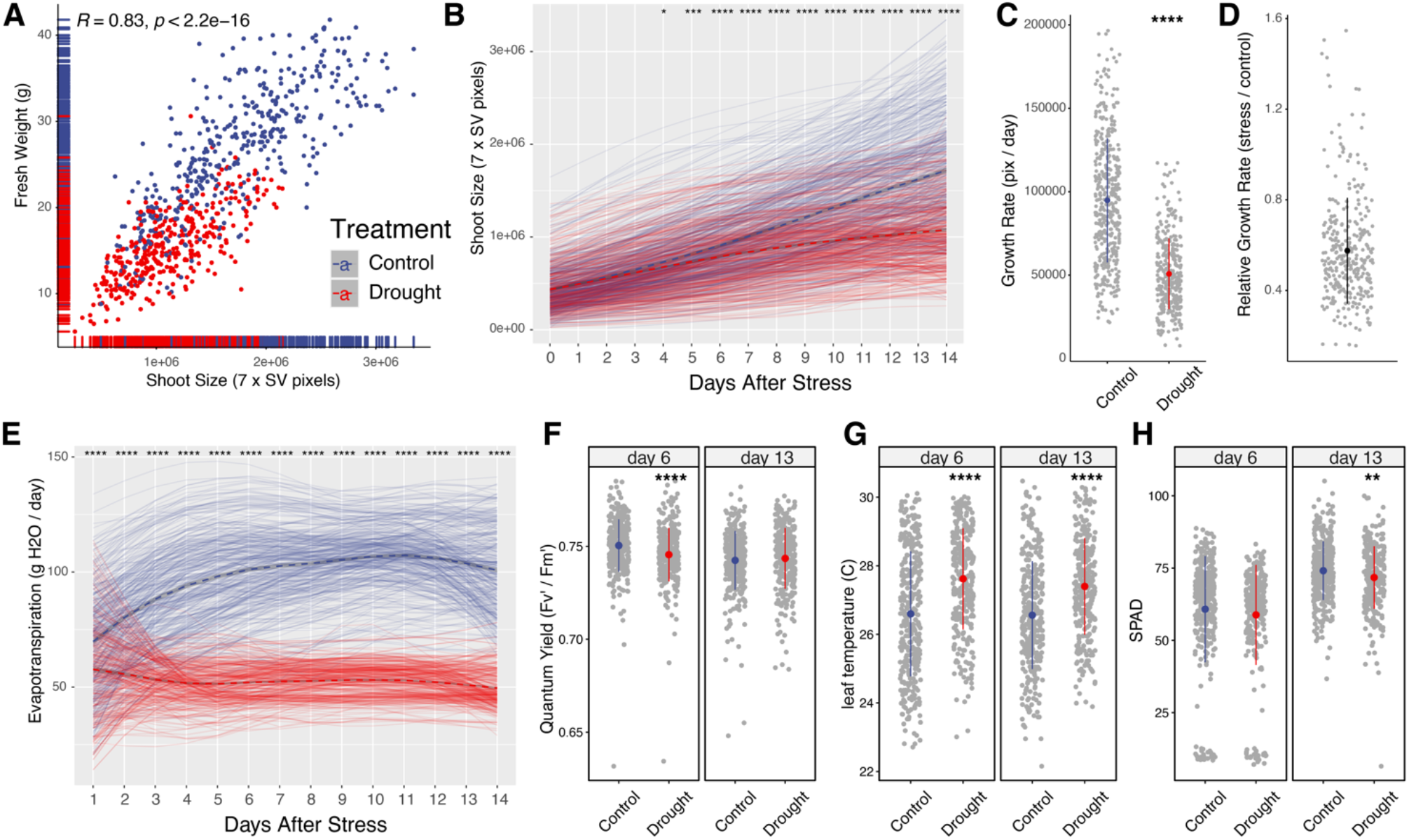
Drought stress-induced changes in natural diversity panel of cowpea. 368 accessions were exposed to 20% soil water-holding capacity (SWHC - Drought) or 60% SWHC (Control) 16 days after sowing. The plants were monitored for increase in shoot biomass and evapotranspiration over the period of two weeks. **(A)** Correlation between digital biomass and fresh weight on the last day of measurement. Pearson correlation coefficient (R) and p-value of the correlation are presented in upper left corner of the graph. **(B)** Increase in digital biomass over the experiment for plants exposed to Control and Drought treatments. **(C)** Growth rate was calculated by fitting linear function to digital biomass for each accession for the duration of the treatment. **(D)** Relative growth rate was calculated for each accession by dividing genotype-specific growth rate recorded under drought condition by growth rate recorded under control conditions. **(E)** Evapotranspiration was estimated for each plant by measuring the pot weight every day of the experiment, watering it to the reference weight, and calculating the difference in weight between consecutive days. The **(F)** Quantum Yield (Fv’/Fm’), **(G)** leaf temperature (C) and **(H)** chlorophyll content were measured using the PhotoSynQ MultiSpeq device. The significant differences between Control and Drought stress were tested using student t-test, and *, **, *** and **** represent p-value < 0.05, 0.01, 0.001 and 0.0001 respectively.

### Identification of new genes underlying drought responses

To identify genetic components underlying the diversity observed in the cowpea miniCore population (**Fig. 7**), we used the collected phenotypic data in combination with the SNPs acquired from SNP array (Muñoz-Amatriaín *et al*., 2021) as input for a Genome-Wide Association Study (GWAS, **Supplemental Table S7**). We examined the identified associations not only for their association strength but also for their predicted effect size (**Fig. 7**). In total, we identified 59 significantly associated SNPs, which could be grouped into 25 loci, based on SNPs falling into 30 kpb window (**Supplemental Table 8**). In total, we identified 10 loci specific to control conditions, 12 drought-specific loci and 3 loci shared between the traits measured under control and drought stress conditions (**Table 1, Supplemental Table 8**). Based on the association strength (-log(p-value) > 5.45), effect size (ß > 3x SD) and the traits, we prioritized 9 loci for further investigation. For all identified associations, we examined the predicted genes in the genome annotation within the linkage disequilibrium (30 kbps) of identified SNPs(Lonardi *et al*., 2019; Muñoz-Amatriaín *et al*., 2021)

**Figure 7.**
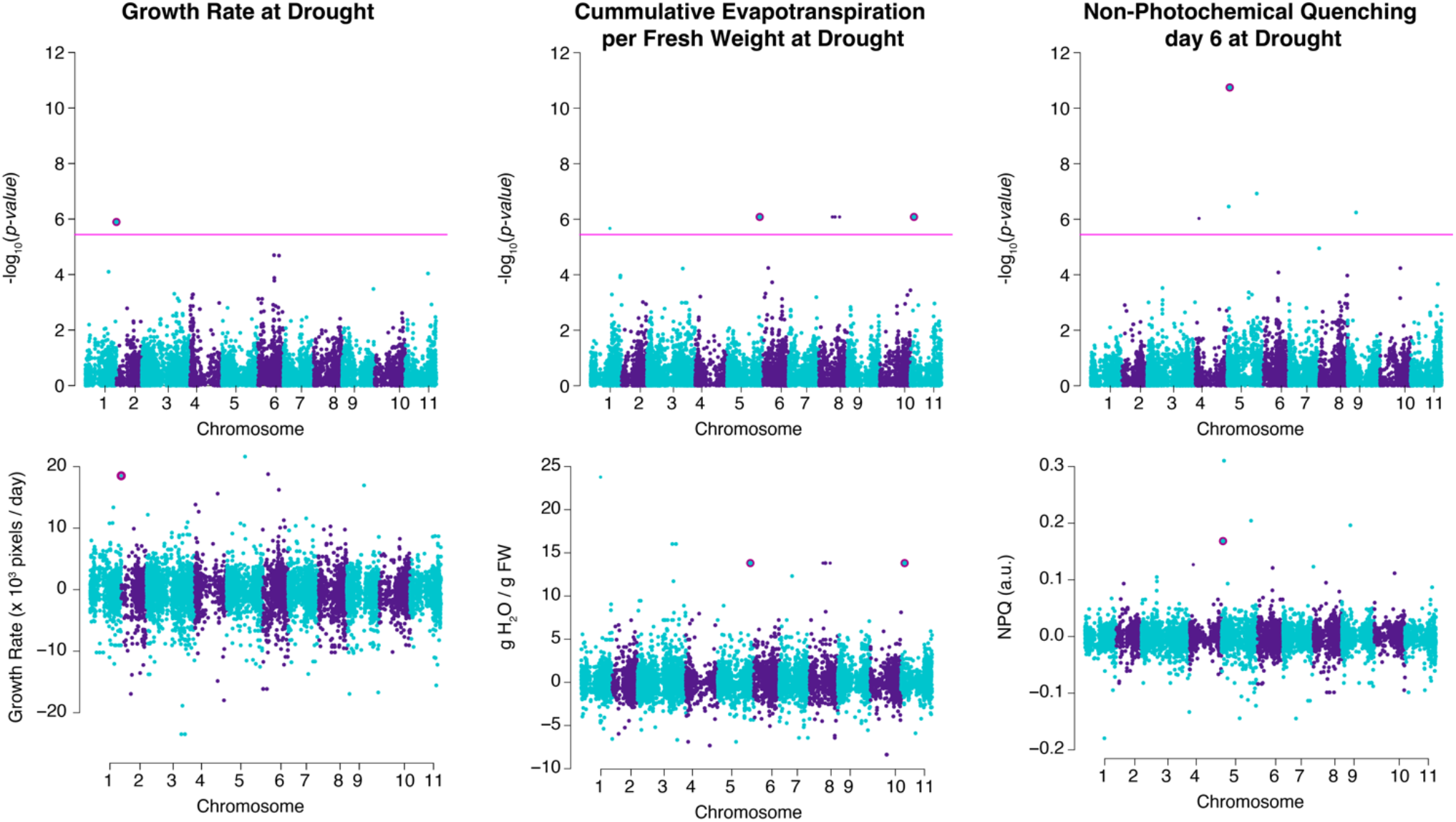
Identification of genetic components of drought stress responses in cowpea through GWAS. Genome Wide Association study was performed using ASReml based script on 368 accessions using 42,711 SNPs and kinship matrix as a co-factor. GWAS associations were examined for overlap between control and drought stress conditions, -log10(p-value) score, minor allele frequency and effect size. The selected associations that were observed exclusively for drought stress treated plants are highlighted in pink. The upper panel represents Manhattan plots, with pink line representing Bonferroni threshold (equal to -log10(0.05 / # SNPs). The bottom panel represents effect size observed for SNPs. The SNPs selected for further investigation (*Table 1*) are highlighted with dark-pink outline.

**Table 1.** GWAS cowpea candidate genes. The summary of the loci with significant association derived from cowpea GWAS panel, including the genomic coordinates (chromosome, location), P-value, beta score, LOD (-log10(p-value) score, and the trait associated with the loci. The candidate genes within linkage disequilibrium of identified SNPs are highlighted in a color scale based on the distance of genes to SNPs. The gene function and respective Arabidopsis homologs are listed, including the coding for the genes referred to within this manuscript (BTI.code). The traits associated with the identified SNPs through GWAS are abbreviated as follows: Growth Rate (GR), Non-photochemical quenching (NPQ), Evapotranspiration rate per seedling fresh weight (EVT.p.FW), whereas Drgth stands for associations identified exclusively under drought stress conditions.

Growth rate under drought stress was associated with one SNP on chromosome 1, positioned within the coding region of Vigun01g250400, which, according to the genome annotation, is a putative homolog of the Arabidopsis gene AT4G14180, which encodes a Putative Recombination initiation Defect protein (AtPRD1), required for DNA double-strand break formation during meiosis. The two genes directly up and downstream of the associated SNP (Vigun01g250500 and Vigun01g250600) are hypothesized to encode pentatricopeptide repeat and zinc-finger (C2H2 type) family proteins, respectively. We identified two drought-specific associations with evapotranspiration use efficiency under drought stress (**Fig. 7**, **Table 1**), located on chromosomes 5 and 8. The association on chromosome 5 was found within Vigun05g246700, which is a putative homolog of Arabidopsis AT3G25830, encoding a monoterpene 1,8-cineole synthase (AtTPS-Cin). The monoterpene 1,8-cineole was previously associated with the decreased root growth in *Brassica campestris* (Koitabashi *et al*., 1997), and AtTPS-Cin is expressed in Arabidopsis roots (Chen *et al*., 2004). The association on chromosome 8 was found within Vigun08g112100, which encodes a putative cowpea homolog of GLY1 (AT2G40690), a nuclear-encoded NAD-dependent glycerol-3-phosphate dehydrogenase family protein associated with flux of fatty acids in the chloroplast (Singh *et al*., 2016). The neighboring genes (Vigun08g112000 and Vigun08g112200) encode homologs of sucrose transporter (SUT4, AT1G09960) and transcription factor (WKRY70, AT3G56400) (**Table 1**).

We found the most significant associations with non-photochemical quenching (NPQ) under drought. However, the majority of these associations (3 out of 5 loci) were also identified under control conditions (**Supplemental Table 7**). The most prominent drought-specific association was located on chromosome 4, within Vigun04g051200, which encodes a cowpea homolog of Arabidopsis glutaredoxin family protein, AT5G39865 (**Table 1**). To evaluate the function of identified genes in drought stress response, we examined the available homozygous T-DNA insertion lines of the putative Arabidopsis homologs of the identified cowpea candidate genes (**Table 1, Supplemental Table 9, Fig. S6-S9**). The available mutants were grown alongside Col-0 wild-type in the soil pots, and at 2 weeks after germination, watered to target weight corresponding to 60 and 10% soil water-holding capacity for control and drought treatments respectively. The plants were imaged every 30 minutes using the PhenoRig system, while their weight was recorded and adjusted every 2nd day using AWWEsmo. The initial screen revealed that out of 43 T-DNA insertion lines, six and two lines developed significantly larger or smaller rosettes, respectively when compared to Col-0 under drought stress conditions (t-test p-value < 0.05, **Fig. S6, S7**). Eight and four T-DNA insertion lines showed respectively increased or decreased evapotranspiration rates under drought stress compared to Col-0 (**Fig. S8**). Nine and two lines showed increased or decreased leaf temperature respectively, whereas five lines showed a significant decrease in non-photochemical quenching (**Fig. S8**). In total, six T-DNA insertion lines showed overlap in the measured phenotypes under drought conditions (EVT2-2, EVT3-2, EVT6-2, EVT8, GR4-1 and N-3 targeting 1,8-cineole synthase, alpha carbonic anhydrase 7, WRKY70, CAAX aminoterminal protease family, xyloglucan endotransglucoselase / hydrolase 16, and pentatricopeptide repeat protein respectively, **Table 1**). As it is possible that other alleles targeting these genes were not detected as significantly different from Col-0 due to a low number of replicates per genotype per condition (n=4), we performed an additional experiment with an increased number of replicates (n=12) (**Fig. 8-10**).

**Figure 8.**
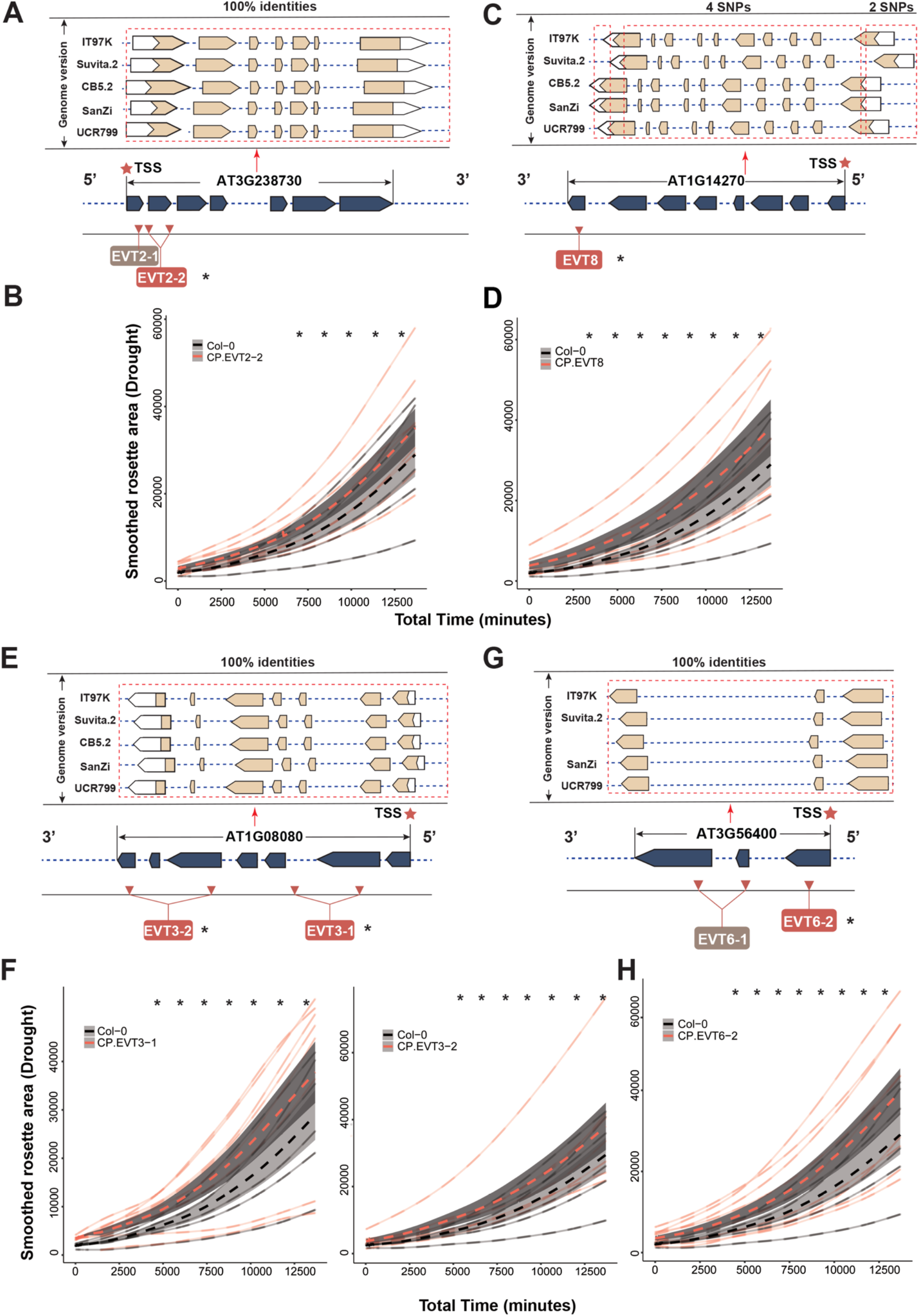
Plant growth of evaporation loci-associated mutants relative to Col-0 plants under drought condition. **(A)** Gene model of 1,8-cineole synthase (AtTPS27, AT3G25830) and location of EVT2 T-DNA insertion. **(B)** Growth of Col-0 and two AtTPS27 mutants under drought condition. **(C)** Gene model of CAAX amino terminal protease (AT1G14270) and location of EVT8 T-DNA insertion **(D)** Growth of Col-0 and protease mutants under drought condition. **(E)** Gene model of alpha carbonic anhydrase 7 (AtACA7, AT1G08080) and the and location of EVT3 T-DNA insertions. **(F)** Growth of Col-0 and two AtACA7 mutants under drought condition. **(G)** Gene model of AtWRKY70 (AT3G56400) and location of EVT6 T-DNA insertions. **(H)** Growth of Col-0 and AtWRKY70 mutant under drought condition. Only insertion sites with significant differences of leaf area under the drought condition were displayed (t-test: P < 0.05)

The 13 selected mutants were grown under control and drought (20% soil-water holding) conditions, except this time the drought stress was imposed using electric fans to ensure that the plants would reach their target weight within the first two days of the stress imposition. Under both control and drought conditions, all 13 of the mutant lines used the same amount of water as the Col-0 plants (**Fig. S9**), including the identified loci initially linked to evapotranspiration (**Fig. 8**). Significantly larger rosettes were observed in the mutant lines targeting genes encoding 1,8-cineole synthase (AtTPS27, EVT2-2), CAAX amino terminal protease (EVT8), Alpha carbonic anhydrase 7 (AtACA7, EVT3-1, EVT3-2) under drought conditions but not under control conditions (**Fig. 8, Fig. S10**). The mutant line targeting WRKY70 (EVT6-2) had significantly (p < 0.05; t-test) larger rosettes under both control and drought conditions (**Fig. 8, Fig. S10**). Under control conditions alone, for the last part of the experiment, we observed significantly larger rosette sizes in EVT6-1, which also targets WRKY70 (**Supplemental fig. 10, 11**). For the CAAX protease and AtACA7, we observed that all tree-screened T-DNA insertion lines showed a significant increase in rosette size (**Fig. 8 D, F**). AtACA7 is predominantly expressed in root stele (Brady et al., 2007), and its expression under abiotic stress was not reported in previous studies (Kilian *et al*., 2007). CAAX protease is expressed in the new leaves (Klepikova *et al*., 2016), and its expression does not change in response to drought or osmotic stress (Kilian *et al*., 2007). Only one of the two screened T-DNA insertion lines for TPS27 and WRKY70 showed a significant increase in rosette size under drought stress (**Fig. 8 B, H, Fig. S11**). Based on previous data, WRKY70 is expressed in the senescing leaf petiole (Klepikova *et al*., 2016), but its expression is unaltered by drought stress in Col-0 (Kilian *et al*., 2007). On the other hand, TPS27 is known to be expressed in the root stele (Brady *et al*., 2007) and is increased by exposure to osmotic, salt, and drought stresses (Kilian *et al*., 2007). These results suggest a new function for the identified candidate genes in drought resilience through maintenance of rosette growth under both control and drought stress conditions.

Additionally, we observed larger (p <0.05; t-test) rosette size in drought and control stressed plants in three (NPQ6-2, NPQ6-4, NPQ6-5) of the mutant lines targeting the Arabidopsis homolog to the gene associated with non-photochemical quenching in cowpea, pentatricopeptide repeat super-family protein (AtPPR, **Fig. 9, Fig. S10**). For the NPQ6-1 mutant line, significantly larger rosette sizes were observed only under drought stress conditions (**Fig. 9**). Within the cowpea pangenome (REF TO PANGENOME), the 5’ UTR is the predominant site of sequence variation, containing one indel mutation and 4 SNPs (**Fig. 9A**). Only one of the five studied insertion lines did not show significant changes in rosette size (NPQ6-3, **Fig. S11**). As the location of this insertion line is beyond the 3’ UTR, it is likely that this mutation does not disturb expression of the gene. PPR expression is ubiquitous (Klepikova *et al*., 2016), and unaltered in response to drought or osmotic stress (Kilian *et al*., 2007). These results suggest an involvement of pentatricopeptide repeat super-family protein in maintenance of rosette growth under control and drought stress conditions.

**Figure 9.**
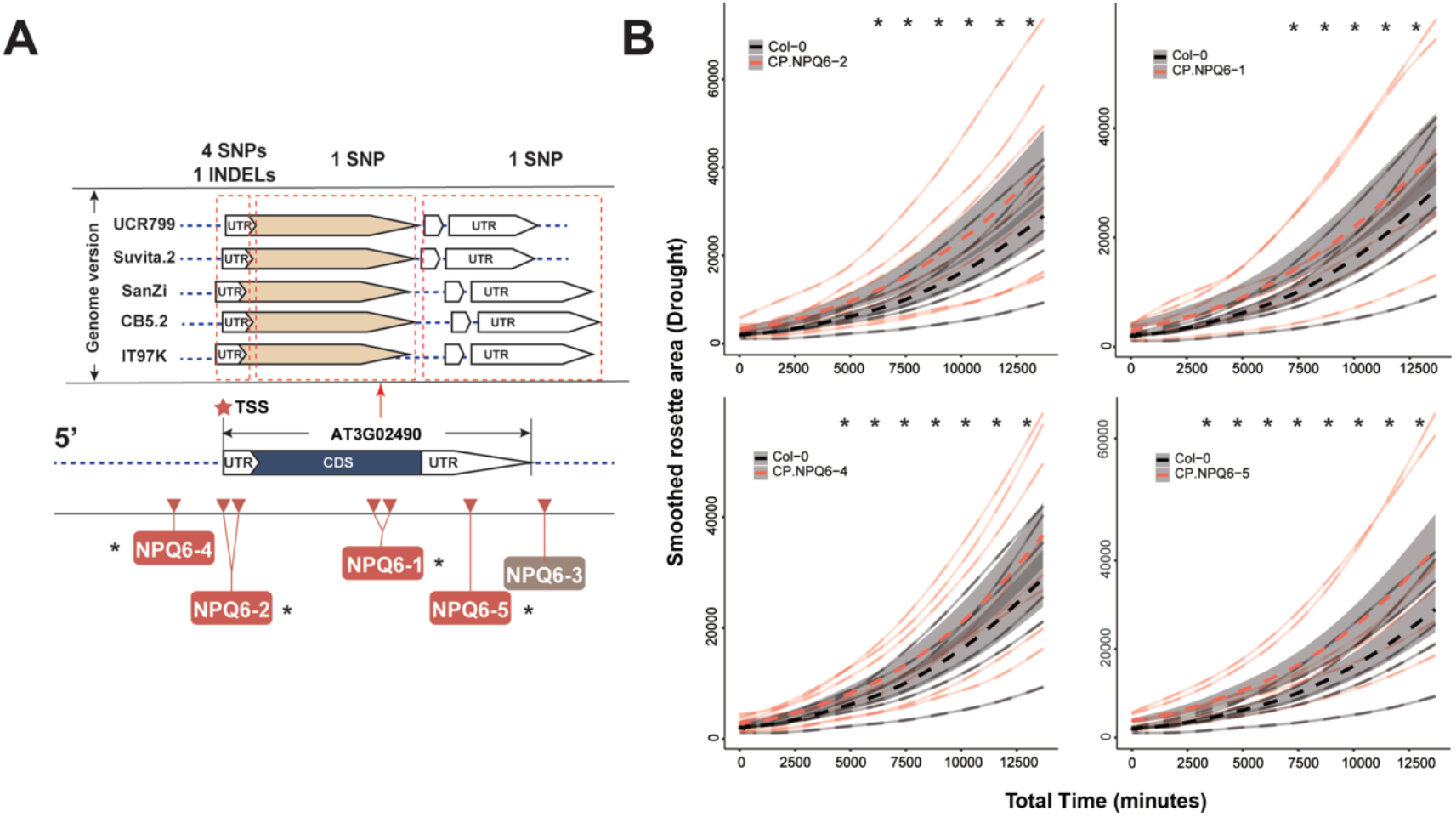
Plant growth of NPQ loci-related mutants relative to Col-0 plants under drought condition (A) Gene model of pentatricopeptide super family protein (AtPPR, AT3G02490) and location of the five T-DNA insertion sites of mutants. In addition, the orthologous gene (Blastn search: E-value < 1 e05, identities > 95%) from cowpea was identified from five published genome assemblies and gene annotations (UCR799, Suvita.2, SanZi, CB5.2, and IT97K). The multiple-alignment of five orthologous genes revealed the high-level conservation across coding region (1 SNPs among five genes), as well as the 3’UTR region (1 SNP). **(B)** Growth of Col-0 and AtPPR mutants under drought condition. Only the four out of five mutants along with significant difference of leaf area between wild-type and mutants under drought condition were plotted (t-test: *p-value* < 0.05)

**Figure 10.**
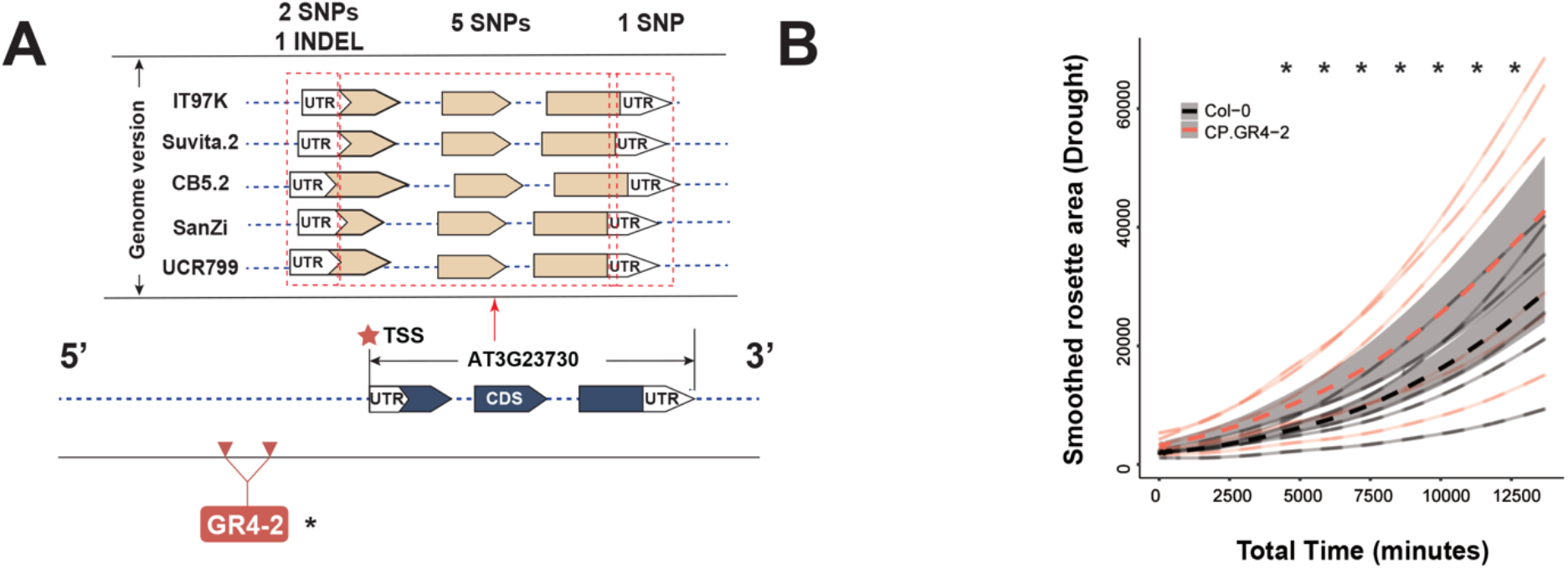
Plant growth of GR loci-related mutants relative to Col-0 plants under drought condition. **(A)** Gene model of xyloglucan endotransglucosylase / transferase 16 (AtXTH16, AT3G23730) which is associated with growth rate and location of the T-DNA insertion site of one mutant. In addition, the orthologous gene from cowpea was identified from five versions of genome assembly and annotation (UCR799, Suvita.2, SanZi, CB5.2, and IT97K). The multiple-alignment of five orthologous genes revealed the different levels of conservation across the 5’UTR (2 SNPs and 1 INDEL), the coding region (5 SNPs),and the 3’UTR (1 SNP). **(B)** Growth of Col-0 and AtXTH16 mutants under drought condition. This loci revealed the significant difference of leaf area in between wild-type and mutant (t-test: *p-value* < 0.05)

In the final locus that we have investigated in further detail, the Arabidopsis mutant line targeting xyloglucan endotransglucosylase/transferase 16 (AtXTH16), which is homologous to the gene associated with growth rate under drought stress in cowpea, we observed larger (p < 0.05; t-test) rosette size under both control and drought conditions (**Fig. 10, Fig. S10**). Within the cowpea pangenome (Liang *et al*., 2022), the majority of the sequence variation resides within the gene coding region (5 SNPS, **Fig. 10A**) and 5’UTR (2 SNPs, 1 indel mutations). Expression of AtXTH16 has been reported to occur predominantly in the developing leaves (Klepikova *et al*., 2016), and is decreased in the initial stage of drought stress exposure (Kilian *et al*., 2007). While we could only find one T-DNA insertion line for this gene, the above results suggest that XTH16 might play a role in rosette growth under control and drought stress conditions. Overall, we have identified six new genes to play a role in drought stress tolerance potentially by maintaining the rosette growth, as increased rosette size was observed in both control and drought stress conditions (**Fig. 8, 9, 10**, Fig. **S-control-growth-rates**)

## Discussion

Plant phenotyping plays a crucial role in plant breeding, genetic research, and agricultural productivity, and is essential for understanding plant growth, development, and responses to the environment. Automated plant phenotyping methods provide valuable information that can be used to improve crop productivity and sustainability, and to address global challenges such as food security and climate change. Implementing computational tools into plant phenotyping enables the efficient processing and analysis of large amounts of images (Fahlgren *et al*., 2015a; Gehan *et al*., 2017). As plant phenotyping methods rely increasingly on computational tools, their adoption is often hindered by the requirement of advanced programming skills and the lack of user-friendly interfaces (Klukas *et al*., 2014; Zhou *et al*., 2021) Most wet labs consist of scientists with limited coding or engineering experience, and thus it is necessary to democratize the phenotyping tools derived from new technologies and algorithms, and to increase the level of accessibility and convenience of novel tools for the plant scientists.

Here, we present a cost-effective all-in-one system, consisting of hardware designs and data analysis pipelines that streamline individual workflows into one experimental framework **(Fig. 3)**. We aim to break down the bottleneck in plant phenotyping by combining phenotypic image collection with data analysis for users without coding experience. The developed tools allow for navigation between plant images, intermediate datasets, and data analysis. By following the detailed instructions, users can collect data and execute data processing and analysis steps without prior knowledge of programming and engineering.

The most significant advantage of the developed system is its low cost and high versatility. As presented in the results, the hardware pieces were constructed with lightweight and low-cost materials that are simple to put together (**Fig. 1** and **S1)**. Moreover, the system’s adaptability to a broad range of research contexts was demonstrated by imaging a wide range of plant species. We demonstrated the precision of the system in predicting digital biomass for species grown under controlled environmental conditions (**Fig. 3** and **4**). Our system has been developed to monitor plant growth during the early vegetative developmental stage, as this early stage has been the focus of many experiments under controlled conditions (Behmann *et al*., 2014; Zhang *et al*., 2020), and RGB image-based phenotyping is reliable in predicting biomass (Matiur *et al*., 2017; Banerjee *et al*., 2020). The other Raspberry Pi-based systems, such as “SPIRO” and “PhenoWell” are also controlled by low-cost computers, offering sustainable phenotyping solutions for high-throughput phenotyping (Ohlsson *et al*., 2021; Li *et al*., 2023). However, these systems are restricted to imaging smaller plants with less complex 2D architectures. By contrast, our system can be expanded in size, allowing for monitoring larger and more complex plants.

To further enable fast and reproducible data analysis, we developed an interactive ShinyApp, RaspiPheno-App, to decode plant genotype and treatment information, as well as to perform the data curation, smoothing, and visualization (https://github.com/Leon-Yu0320/BTI-Plant-phenotyping/tree/main/RasPiPheno_APP). This approach could potentially reduce the downstream workload compared to some other systems which have no specific data analysis suite included (Czedik-Eysenberg *et al*., 2018; Feldman *et al*., 2021). Similar to other ShinyApps for biological data analysis (Julkowska *et al*., 2019; Ge, 2020; Xiao & Lam, 2022), the RaspiPheno-App provides a fast and straightforward approach for data processing. Moreover, RaspiPheno-App offers compatibility with data input derived from previous steps, as it is fine-tuned to match the data output format from the RaspiPheno pipeline. This direct data adoption requires no further data reformatting, reducing the learning curve required for new users. Integration of the ShinyApp into the suite of developed tools allows for decoding of information and assigning plant genotype and/or treatment to individual plants, as well as other meta-features of an experimental design. As a result, the streamlined ShinyApp enables the acquisition of high-quality plots and systematic statistical analysis of plant growth, to address the specific biological questions about the physiological and genetic basis of plants.

Within this work, we have demonstrated the system’s usability using a cowpea diversity panel to drive the discovery of new genes involved in environmental resilience. The observed phenotypic variation in cowpea (**Fig. 6**) was used as an input for GWAS (**Fig. 7**) to identify new genetic loci contributing to drought resilience. Previous studies have identified cowpea as an important donor for drought tolerance traits (Muñoz-Amatriaín *et al*., 2021), however cowpea phenotyping has thus far not been performed on dynamic traits, such as growth rate or evapotranspiration during drought exposure (Muchero *et al*., 2008, 2009; Verbree *et al*., 2015; Ravelombola *et al*., 2020; Nkomo *et al*., 2022). We observed that, similarly to other crop plants (Yuan & Liu, 2012), cowpea reduces its evapotranspiration (**Fig. 6 E**). On the other hand, the photosynthetic efficiency was only affected at the early phase of drought stress exposure (**Fig. 6 F**). It has been previously reported that Arabidopsis plants under salt stress have been found to decrease quantum yield during early salt stress exposure, and that this impacted growth maintenance (Awlia *et al*., 2021).

To explore the potential genes involved in drought resilience in cowpea, we examined available T-DNA insertion mutants in Arabidopsis homologs to the set of cowpea genes with significant trait-associations. Our experiments revealed new genes potentially involved in drought resilience (**Fig. 8, 9 and 10**), highlighting the usefulness of these low-cost, large scale, phenotyping approaches. Not all studied T-DNA insertion lines showed significant changes in rosette growth rate under drought conditions, and thus require further validation (e.g. AT3G25830 encoding 1,8-cineole synthase, AT1G14270 encoding CAAX amino terminal protease, AT3G56400 encoding AtWRKY70, and AT3G23730 encoding xyloglucan endotransglucosylase / transferase 16, **Fig. 8 A-D and G-H**, **Fig. 10**). Two out of two studied T-DNA insertion lines targeting alpha carbonic anhydrase 7 (AtACA7, AT1G08080) and four out of five T-DNA insertion lines targeting pentatricopeptide repeat super-family protein (AtPPR, AT3G02490) showed convincing evidence for its involvement in drought resilience (**Fig. 8 E-F** and **Fig. 9** respectively). Carbonic anhydrases are an abundant protein family that has multiple isoforms and acts in carbon assimilation and photosynthesis (Momayyezi *et al*., 2020). The three families, alpha, beta, and gamma, are considered to have evolved separately (Momayyezi *et al*., 2020) but have similar functions (Moroney *et al*., 2001). The alpha carbonic anhydrases have the overall lowest expression level (Polishchuk, 2021). While no previous reports cover the role of alpha carbonic anhydrases in drought stress resilience, the beta family has been studied extensively. The expression of beta carbonic anhydrases has been shown to both increase (Polishchuk, 2021) and decrease (Wang *et al*., 2016; Han *et al*., 2019; Momayyezi *et al*., 2020) under drought. These contrasting findings, as well as our results, indicate the need for further investigation into the role of alpha carbonic anhydrases in drought response in plants.

Pentatricopeptide repeat proteins (PPRs) in plants are known for their wide range of molecular functions, including photosynthesis and environmental stress responses including drought stress (Barkan & Small, 2014). Arabidopsis plants exhibited improved growth performance under drought stress when the expression of the PPR protein SLG1 was disrupted (Yuan & Liu, 2012). However, Arabidopsis lines overexpressing another PPR protein, SOAR1, also performed better under drought stress compared to wild type (Jiang *et al*., 2015). To our knowledge, the PPR identified in this study (AT3G02490) has not been previously studied in detail or reported as contributing to drought resilience. These findings highlight the diversity and complexity of PPR proteins, emphasizing the need to characterize our identified PPR further. The molecular context of these genes will be the focus of future studies, revealing new mechanisms of drought resilience across a wider range of species. Our results illustrate the potential of the developed setup in gene discovery and identification of resilience mechanisms for a wide diversity of plants. In the future, the identified genes can serve as attractive targets for breeding or genetic modification to further contribute to crop stress resilience and food security.

Image-based phenotyping is the workhorse of high-throughput phenotyping (Langstroff *et al*., 2022; Hall *et al*., 2022). The application of high-throughput phenotyping in simple and cost-efficient systems, like the one described in our manuscript, carries the potential for a broader impact to plant science research without the prohibitively high costs (Yang *et al*., 2013; Zhou *et al*., 2018; Du *et al*., 2021). The wide adoption of cost-effective solutions can lead to tremendous progress in studying stress responses and identifying new genetic components of environmental resilience.

## Acknowledgments

We would like to thank the BTI workshop team, Lucas Burke and Don Slocum, for their help in setting up the wooden frames for the PhenoRigs and PhenoCages. We would like to acknowledge Ms. Holly Carter and Ms. Rose Deshler (2021 REU students) for their help in troubleshooting data acquisition on the PhenoRigs. The authors would also like to acknowledge Dr. Maria Munoz-Amatrain for sharing the population of the cowpea miniCORE diversity panel. The work is supported by USDA-NIFA grant number 2022-67013-36212. The authors would like to acknowledge NSF-IOS 2023310 and 2102120 (to ADLN). The work on developing affordable phenotyping systems has been supported through Goelet Foundation.

## Supplemental Material Legends

### Supplementary Figures

**Figure S1. Components of PhenoRig and PhenoCage system.** The PhenoRig and PhenoCage were constructed with sized pieces and accessories. **(A)** Illustration of different components used for Phenorig main-body construction, including wood bars in five different sizes (Parameters: Supplemental table-S1) **(B)** Indication of image collection accessories of PhenoRig, including two cameras and connection ribbons **(C)** Illustration of different components used for PhenoRig main-body construction, including wood bars in seven different sizes (Parameters: Supplemental table-S2) **(D)** Indication of illumination accessories, including three LED lights. **(E)** 3-D printed accessories for holding cameras and LCD touchscreen panel

**Figure S2. 3-D printed accessories used for phenotyping system (A)**The LCD touch screen holder for PhenoRig **(B)** LCD touch screen holder for PhenoCage **(C)** Raspi camera and Raspi computer holders for PhenoRig system **(D)** Trellis used for plant individual support during plant growth **(E)** Raspi camera and Raspi computer holders for PhenoCage system

**Figure S3. Construction of the imaging trellis for tepary beans.** In order to provide resources for the phenotyping of the climbing and prostrate plants, we developed an imaging trellis. **(A)** The trellis was designed using TinkerCAD and 3D printed using a transparent PLA filament. The trellis is designed so that two trellises can be stacked upon each other providing approximately 30 cm of vertical support for the climbing plant. **(B)** An image of the tepary bean plant with two trellises providing vertical support. The trellis is merging with the white background of the PhenoCage and is barely detectable with an eye. **(C)** The same image including the imaging trellis analyzed using PlantCV for the projected shoot area.

**Figure S4. Drought-induced changes in evapotranspiration of cowpea and tepary beans. (A)** The five cowpea accessions and **(B)** two tepary bean accessions were germinated in soil for 17 days and subsequently, the pots were watered to target weight corresponding to 60% and 10% of soil water-holding capacity for Control and Drought treatment respectively. The pots were measured and watered daily. The difference between pot weight between the watering was used as plant evapotranspiration. The graphs represent the differences in evapotranspiration between the treatments, and the change in evapotranspiration within each pot is represented with a transparent line. The average of all included genotypes is represented with a dashed line. The standard error is represented with a gray ribbon for each treatment. The significant differences between the treatments were calculated using t-test and annotated with *, **, ***, and **** for p-value below 0.05, 0.01, 0.001, and 0.0001 respectively.

**Figure S5. The effects of drought on evapotranspiration in cowpea. (A)** The median evapotranspiration was calculated for each plant over the course of 14 days of stress imposition. **(B)** The cumulative evapotranspiration was calculated by adding the evapotranspiration of each plant over the course of the experiment. **(C)** The Stress Tolerance Index (STI) was calculated by dividing the genotype-specific values for cumulative evapotranspiration observed under drought stress conditions over the cumulative evapotranspiration observed under control conditions. **(D)** The Evapotranspiration Use Efficiency was calculated by dividing an cumulative evapotranspiration over the fresh weight recorded for each plant included in the experiment. The differences between Control and Drought stress were tested using t-test, and *, **, *** and **** indicate p-values below 0.05, 0.01, 0.001 and 0.0001.

**Figure S6. First experimental batch of drought-induced changes in daily growth rate among Arabidopsis T-DNA insertion lines.** The Arabidopsis T-DNA insertion lines (*Table S9*) were germinated on agar plates alongside Col-0, and transplanted to soil 1 week after germination. Every 2nd day the pots were weighed and watered to target weight corresponding to 60 and 10% of soil water-holding capacity for control and drought stress treatment respectively. The daily growth rate (DGR) was calculated over 16 h for each individual day of the experiment, and compared to Col-0 for plants grown under **(A)** Control or **(B)** Drought stress conditions. The transparent lines represent the data recorded of individual replicates, while the genotype-mean and standard error are represented by the dashed line and grey ribbon respectively. The significant differences between Col-0 and each T-DNA insertion line were evaluated using t-test, and *, **, *** and **** signify p-values below 0.05, 0.01, 0.001, and 0.0001 respectively.

**Figure S7. Second experimental batch of drought-induced changes in daily growth rate among Arabidopsis T-DNA insertion lines.** The Arabidopsis T-DNA insertion lines (*Table S9*) were germinated on agar plates alongside Col-0, and transplanted to soil 1 week after germination. Every 2nd day the pots were weighted and watered to target weight corresponding to 60 and 10% of soil water-holding capacity for control and drought stress treatment respectively. The daily growth rate (DGR) was calculated over 16 h for each individual day of the experiment, and compared to Col-0 for plants grown under **(A)** Control or **(B)** drought stress conditions. The transparent lines represent the data recorded of individual replicates, while the genotype-mean and standard error are represented by the dashed line and grey ribbon respectively. The significant differences between Col-0 and each T-DNA insertion line were evaluated using a t-test, and *, **, ***, and **** signify p-values below 0.05, 0.01, 0.001, and 0.0001 respectively.

**Figure S8. Drought-induced changes in evapotranspiration, leaf temperature and non-photochemical quenching among Arabidopsis T-DNA insertion lines.** The Arabidopsis T-DNA insertion lines (*Table S9*) were germinated on agar plates alongside Col-0, and transplanted to soil 1 week after germination. Every 2nd day the pots were weighed and watered to target weight corresponding to 60 and 10% of soil water-holding capacity for control and drought stress treatments respectively. **(A)** The median evapotranspiration was calculated per plant over the course of the entire experiment (2 weeks), while **(B)** leaf temperature and **(C-D)** non-photochemical quenching (NPQ) were evaluated at the last day of the experiment (2 weeks after treatment imposition). The individual points represent the data recorded of individual replicates, while the genotype-mean is represented by the horizontal line. The significant differences between Col-0 and each T-DNA insertion line were evaluated using t-test, and *, **, ***, and **** signify p-values below 0.05, 0.01, 0.001, and 0.0001 respectively.

**Figure S9. Evapotranspiration of mutants versus Col-0.** Evapotranspiration was tracked by recording the amount of water needed to reach target weights every second day for two weeks. (**A)** compares each mutant to Col-0 while under control conditions (60% soil water-holding capacity), whereas **(B)** compares each mutant to Col-0 while under drought conditions (20% soil water-holding capacity). A t-test was performed to identify significant differences between Col-0 and each mutant. No significant differences in evapotranspiration were found between the mutants and Col-0 under both control and drought at any of the studied time points. The color coding for genotypes is reported below each graph. Each transparent line represents the evapotranspiration of one individual plant, and dashed lines and shaded areas represent the genotype average and standard deviation over time respectively.

**Figure S10. The growth rate of studied Arabidopsis mutant lines under non-stress conditions.** The mutants target **(A)** 1,8-cineole synthase (EVT2) **(B)** alpha carbonic anhydrase 7 (EVT3) **(C)** WRKY70 (EVT6) **(D)** CAAX amino terminal protease family protein (EVT8) **(E)** xyloglucan endotransglucosylase/hydrolase 16 (GR4) and **(F)** Pentatricopeptide repeat (PPR) superfamily protein (NPQ6). The list of specific mutants can be found in **Supplemental Table S9.** Growth of Col-0 and individual mutant lines recorded under control conditions (i.e. 60% soil water-holding capacity). Transparent lines represent the growth of individual lines, whereas dashed lines represent the genotype average. The shaded surfaces represent the standard error. Significant differences between wild-type and mutant are indicated with *, **, ***, and **** for t-test p-values < 0.05, 0.01, 0.001, and 0.0001 respectively.

**Figure S11. The growth rate of studied Arabidopsis mutant lines under drought conditions.** The mutant targets including EVT2-1, NPQ6-3, and EVT6-1 were compared to the growth of Col-0 under drought conditions. Transparent lines represent the growth of individual lines, whereas dashed lines represent the genotype average. The pairwise comparison revealed no significant differences between Col-0 relative to each mutant line.

## Supplementary Tables

**Table S1. PhenoRig construction materials** The unit description and quantities of each component used for PhenoRig system construction were listed, along with the hyperlink of purchases.

**Table S2. PhenoCage construction materials** The unit description and quantities for each component used for Phenocage system construction were listed, along with the hyperlink of purchases.

**Table S3. AWEsmo construction materials** The unit description, quantities, and estimated cost for each component used for AAWSome system construction were listed, along with the hyperlink of purchases.

**Table S4 Description and link of R notebooks** The R notebooks of data analysis for each experiment were stored on Rpubs.

**Table S5. Cowpea raw data accessibility link.** The raw data of cowpea GWAS were archived on Zenodo with respective links provided

**Table.S6 Mean Genotype Growth Rates for each condition.** The Growth rates were calculated by fitting a linear function over the increase in the shoot size for each plant, and then the genotype and condition mean growth rate was calculated over available replicates. The Stress Tolerance Index (STI) was calculated by dividing the growth rate calculated for drought conditions over the growth rate calculated for control conditions.

**Table S7. The genotypic means for each accessions used as input for GWAS.** The growth rate (GR) was calculated for each condition over the course of 14 days of experiment. Fresh Weigh (FW) and Dry Weight (DW) was collected at the end of each experiment. Quantum yield under light-adapted conditions (Fv’/Fm’ or FvP_over_FmP), Nonphotochemical quenching (NPQ), chlorophyll content (SPAD), leaf temperature and leaf thickness were collected 6 and 13 days after treatment initiation. Evapotranspiration (EVT) was calculated based on daily EVT values as either the median EVT per plant (EVT.med) or cumulative (EVT.sum) over the whole experiment.

**Table S8. Associations in cowpea miniCORE population identified through GWAS.** The associations identified using ASReml GWAS are listed including the SNP position, count of reference / non-reference allele (AC_1 and AC_0 respectively), minor allele count (MAC) and frequency (MAF). The p-values for the association strength with individual phenotypes (trait) were calculated using EMMA-X based model. The predicted effect size (beta) of the association was calculated using the ASReml script. The distance between individual SNPs was used to group the individually associated SNPs into loci, based on the average linkage disequilibrium in cowpea.

**Table S9. Arabidopsis mutants selection.** The associations identified through GWAS (Table S8) were inspected further for gene coding regions and Arabidopsis homologs within the linkage disequilibrium of each identified locus. The genes within which the SNP was identified are colored red, whereas increasing distance from the SNP is indicated with fading hues of orange. The identified Arabidopsis homologs (based on cowpea genome annotation file) are listed for each gene, if Arabidopsis homolog was identified. The identified Arabidopsis T-DNA insertion lines are listed for each Arabidopsis homolog, and their nomenclature (BTI.coding) as listed within this manuscript. The traits associated with the identified SNPs through GWAS are abbreviated as follow: Growth Rate (GR), Non-photochemical quenching (NPQ), Evapotranspiration rate per seedling fresh weight (EVT.p.FW), whereas Drgth stands for associations identified exclusively under drought stress conditions.

