## Supplemental Figures for "Development of a mobile, high-throughput, and low-cost image-based plant growth phenotyping system"

**Supplementary Figures**

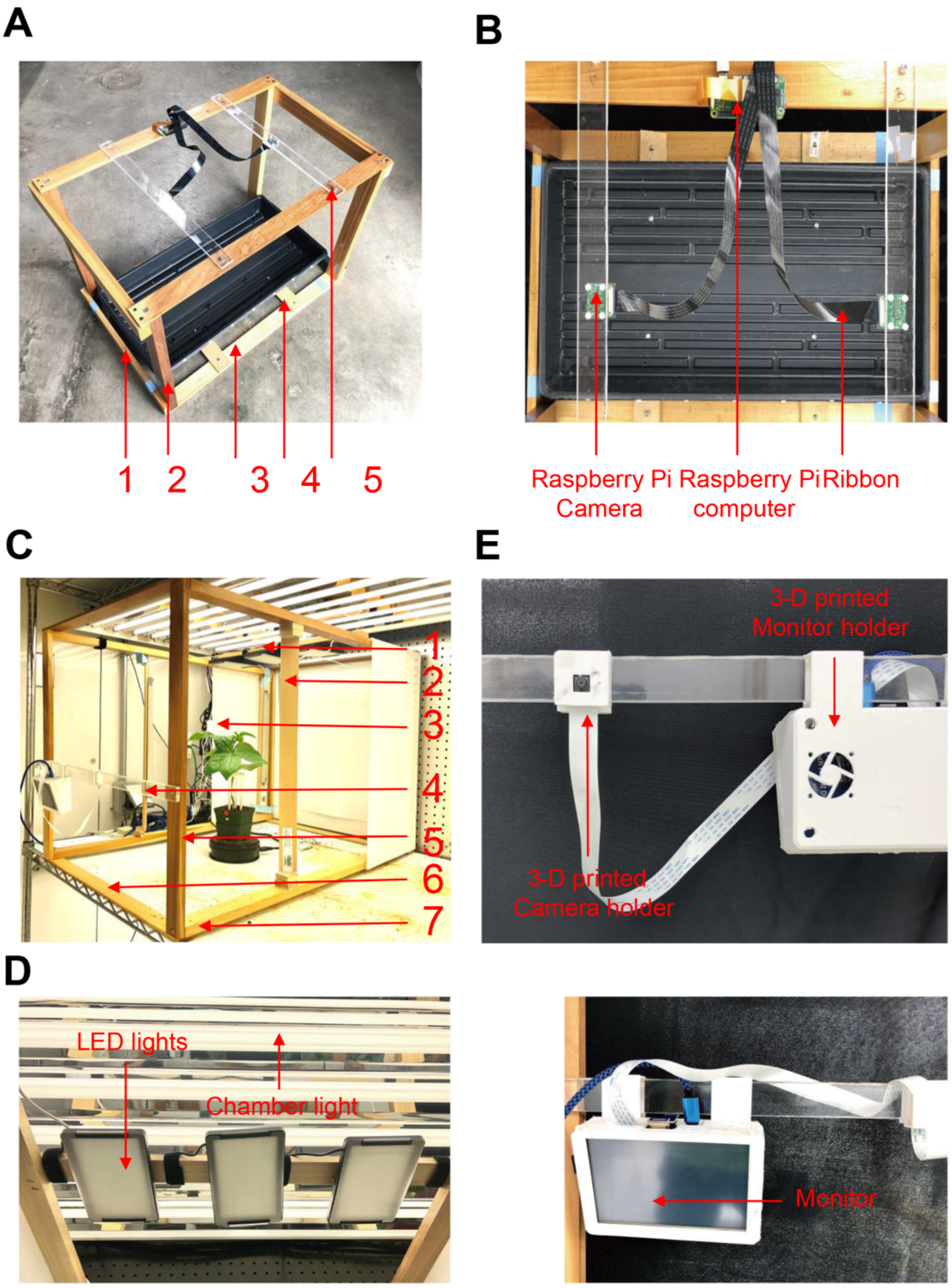


**Figure S1. Components of PhenoRig and PhenoCage system.** The PhenoRig and PhenoCage were constructed with sized pieces and accessories. **(A)** Illustration of different components used for Phenorig main-body construction, including wood bars in five different sizes (Parameters: Supplemental table-S1) **(B)** Indication of image collection accessories of PhenoRig, including two cameras and connection ribbons **(C)** Illustration of different components used for PhenoRig main-body construction, including wood bars in seven different sizes (Parameters: Supplemental table-S2) **(D)** Indication of illumination accessories, including three LED lights. **(E)** 3-D printed accessories for holding cameras and LCD touchscreen panel

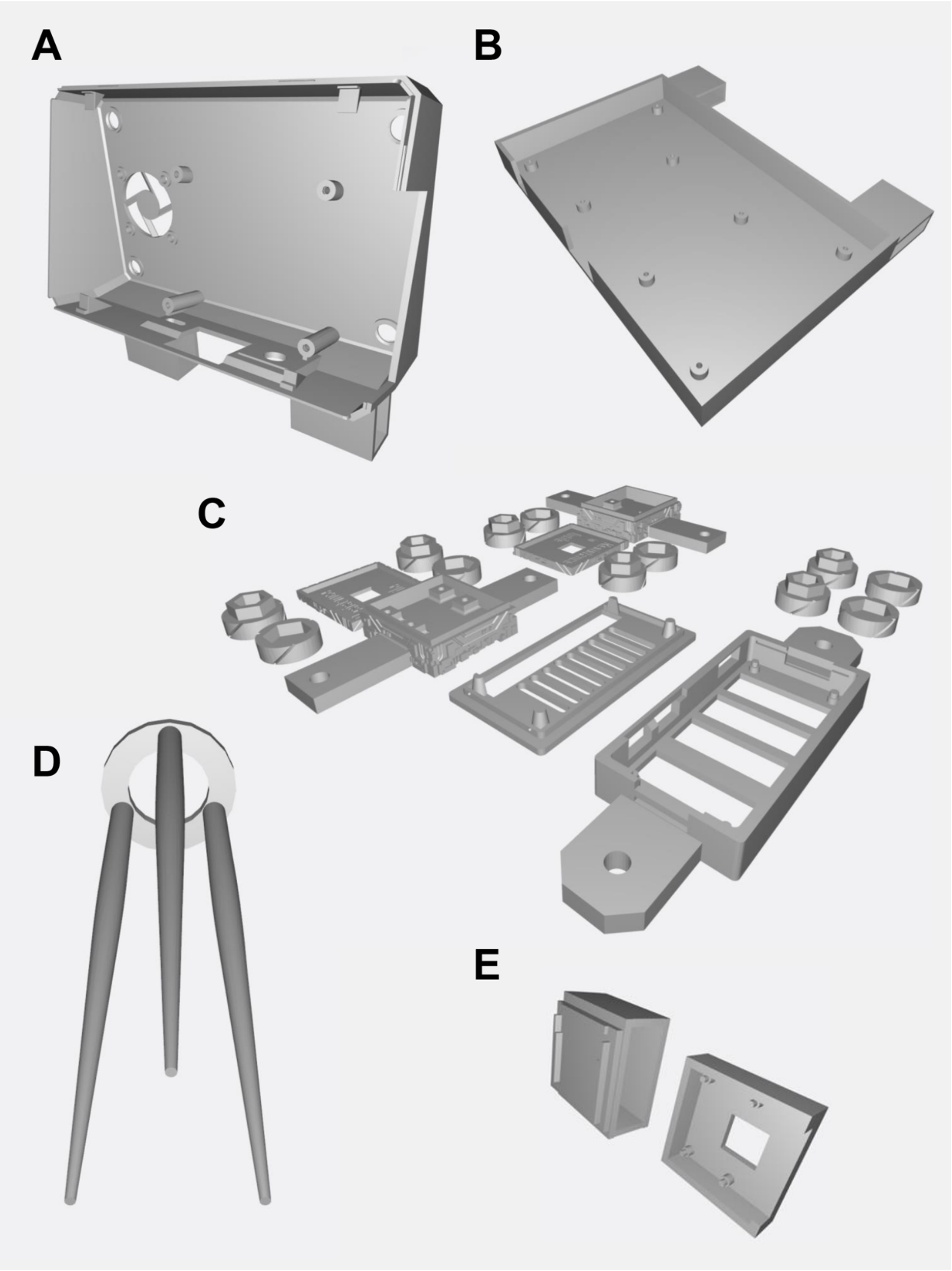


**Figure S2. 3-D printed accessories used for phenotyping system（A）** The LCD touch screen holder for PhenoRig **(B)** LCD touch screen holder for PhenoCage **(C)** Raspi camera and Raspi computer holders for PhenoRig system **(D)** Trellis used for plant individual support during plant growth **(E)** Raspi camera and Raspi computer holders for PhenoCage system

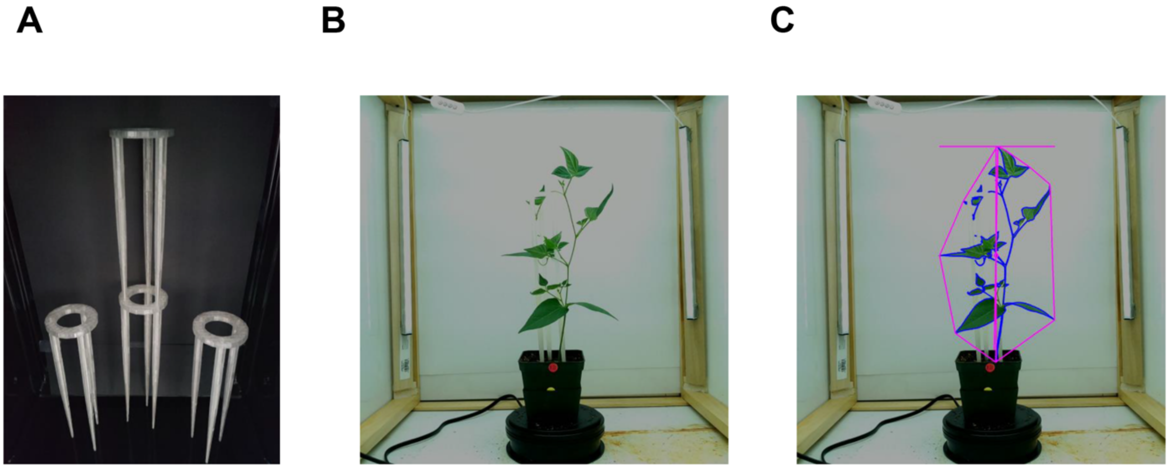


**Figure S3. Construction of the imaging trellis for tepary beans.** In order to provide resources for the phenotyping of the climbing and prostrate plants, we developed an imaging trellis. **(A)** The trellis was designed using TinkerCAD and 3D printed using a transparent PLA filament. The trellis is designed so that two trellises can be stacked upon each other providing approximately 30 cm of vertical support for the climbing plant. **(B)** An image of the tepary bean plant with two trellises providing vertical support. The trellis is merging with the white background of the PhenoCage and is barely detectable with an eye. **(C)** The same image including the imaging trellis analyzed using PlantCV for the projected shoot area.

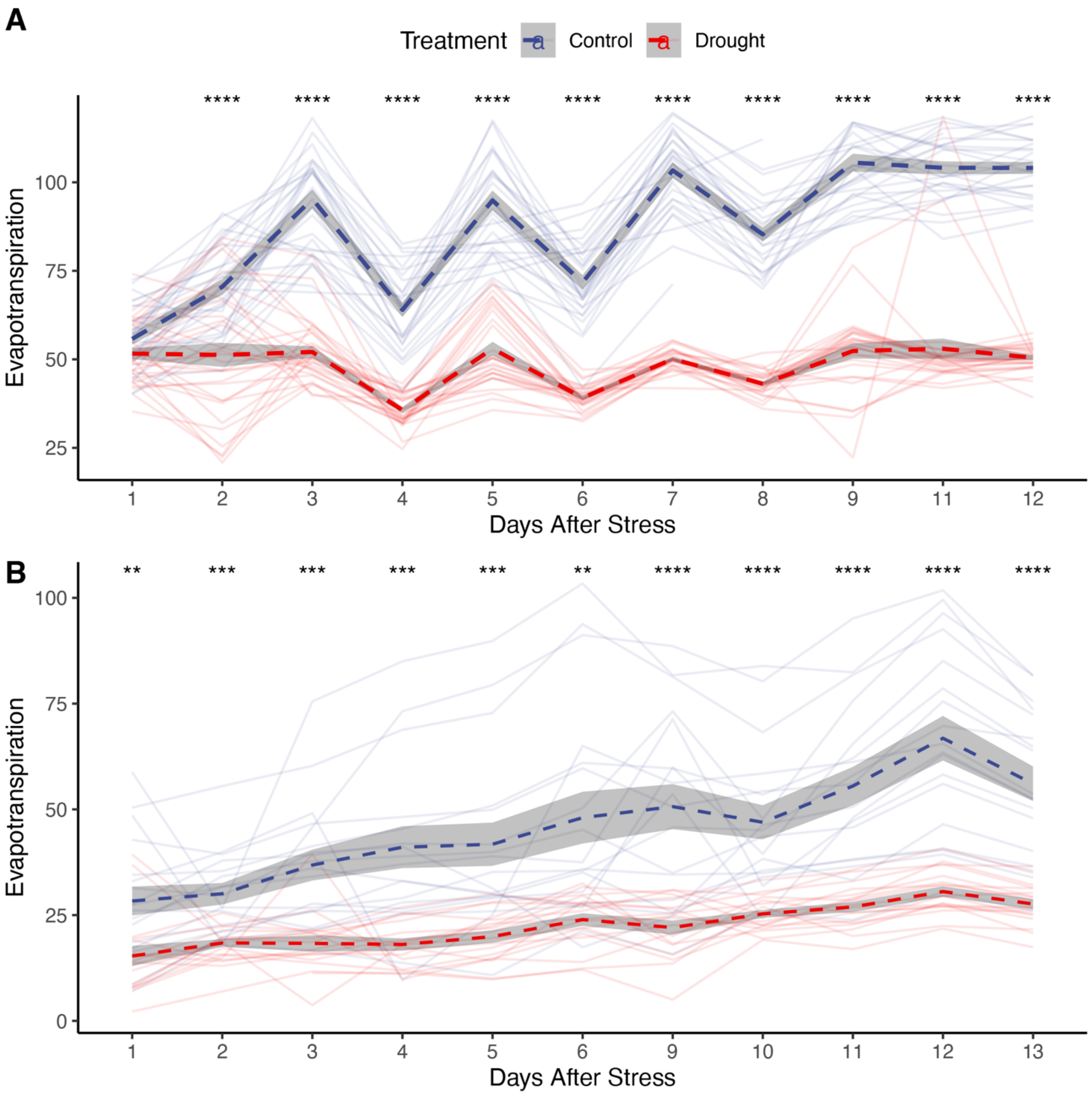


**Figure S4. Drought-induced changes in evapotranspiration of cowpea and tepary beans. (A)** The five cowpea accessions and **(B)** two tepary bean accessions were germinated in soil for 17 days and subsequently, the pots were watered to target weight corresponding to 60% and 10% of soil water-holding capacity for Control and Drought treatment respectively. The pots were measured and watered daily. The difference between pot weight between the watering was used as plant evapotranspiration. The graphs represent the differences in evapotranspiration between the treatments, and the change in evapotranspiration within each pot is represented with a transparent line. The average of all included genotypes is represented with a dashed line. The standard error is represented with a gray ribbon for each treatment. The significant differences between the treatments were calculated using t-test and annotated with *, **, ***, and **** for p-value below 0.05, 0.01, 0.001, and 0.0001 respectively.

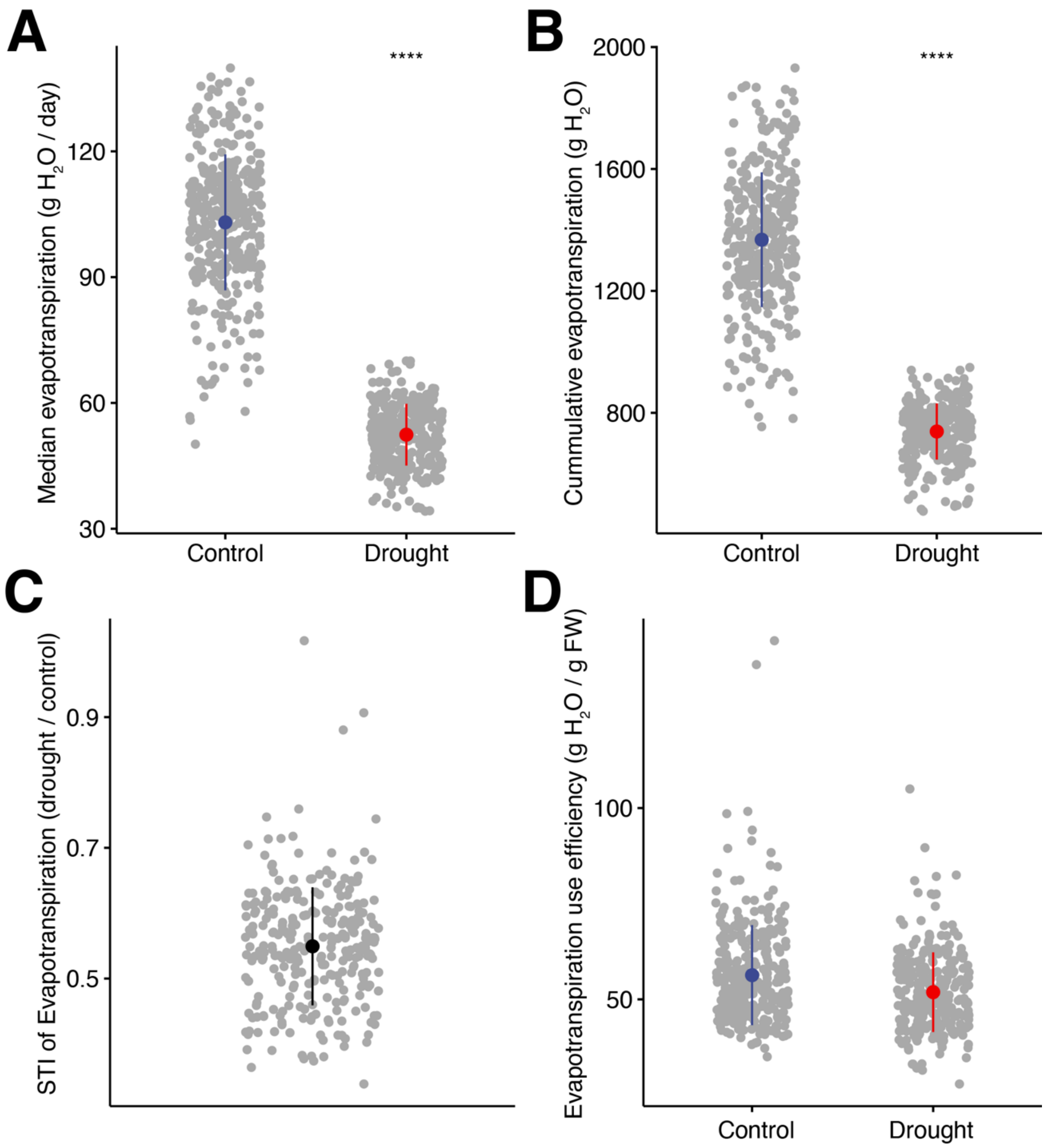


**Figure S5. The effects of drought on evapotranspiration in cowpea. (A)** The median evapotranspiration was calculated for each plant over the course of 14 days of stress imposition. **(B)** The cumulative evapotranspiration was calculated by adding the evapotranspiration of each plant over the course of the experiment. **(C)** The Stress Tolerance Index (STI) was calculated by dividing the genotype-specific values for cumulative evapotranspiration observed under drought stress conditions over the cumulative evapotranspiration observed under control conditions. **(D)** The Evapotranspiration Use Efficiency was calculated by dividing an cumulative evapotranspiration over the fresh weight recorded for each plant included in the experiment. The differences between Control and Drought stress were tested using t-test, and *, **, *** and **** indicate p-values below 0.05, 0.01, 0.001 and 0.0001.

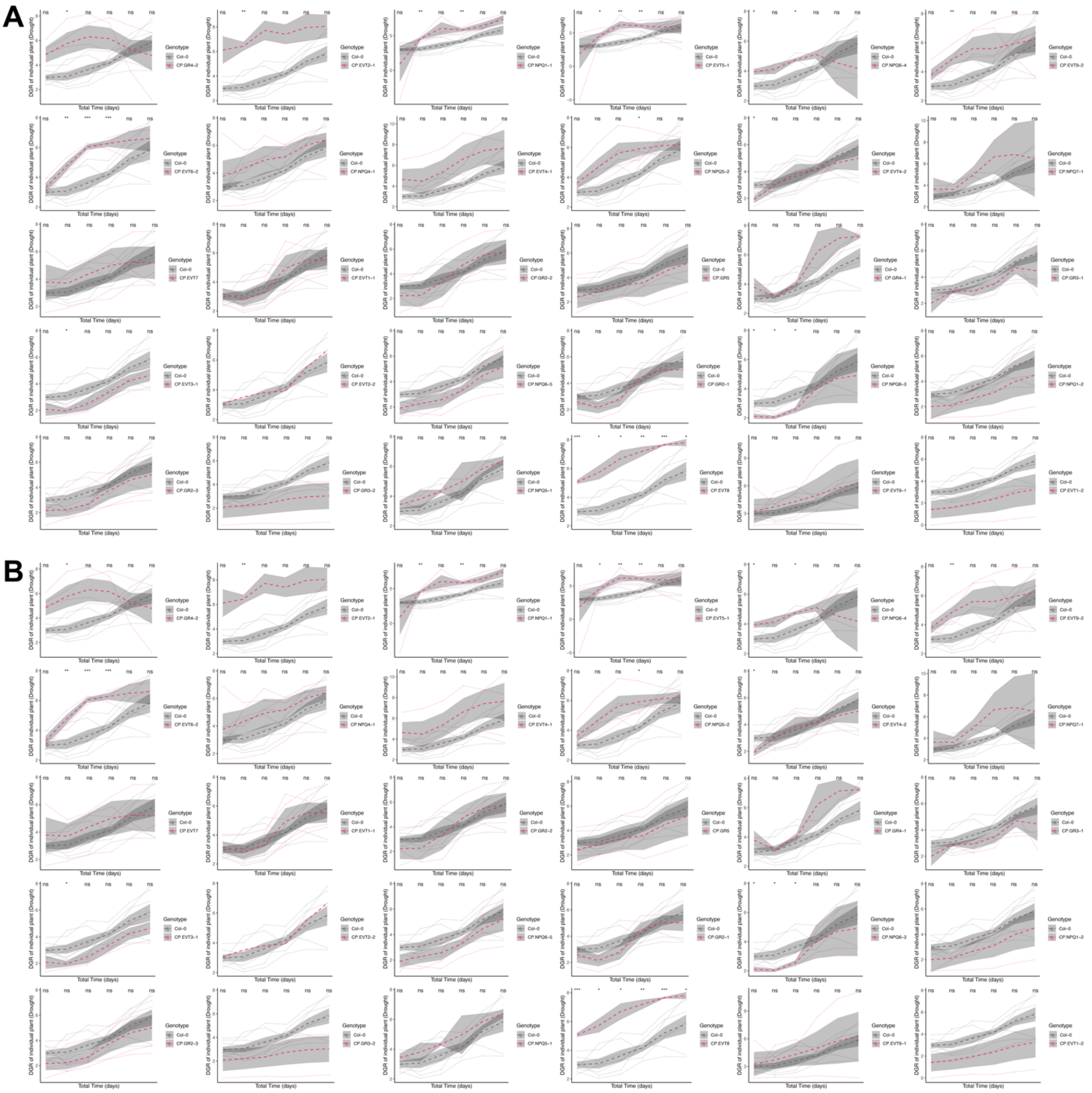


**Figure S6. First experimental batch of drought-induced changes in daily growth rate among Arabidopsis T-DNA insertion lines.** The Arabidopsis T-DNA insertion lines (*Table S9*) were germinated on agar plates alongside Col-0, and transplanted to soil 1 week after germination. Every 2nd day the pots were weighed and watered to target weight corresponding to 60 and 10% of soil water-holding capacity for control and drought stress treatment respectively. The daily growth rate (DGR) was calculated over 16 h for each individual day of the experiment, and compared to Col-0 for plants grown under **(A)** Control or **(B)** Drought stress conditions. The transparent lines represent the data recorded of individual replicates, while the genotype-mean and standard error are represented by the dashed line and grey ribbon respectively. The significant differences between Col-0 and each T-DNA insertion line were evaluated using t-test, and *, **, *** and **** signify p-values below 0.05, 0.01, 0.001, and 0.0001 respectively.

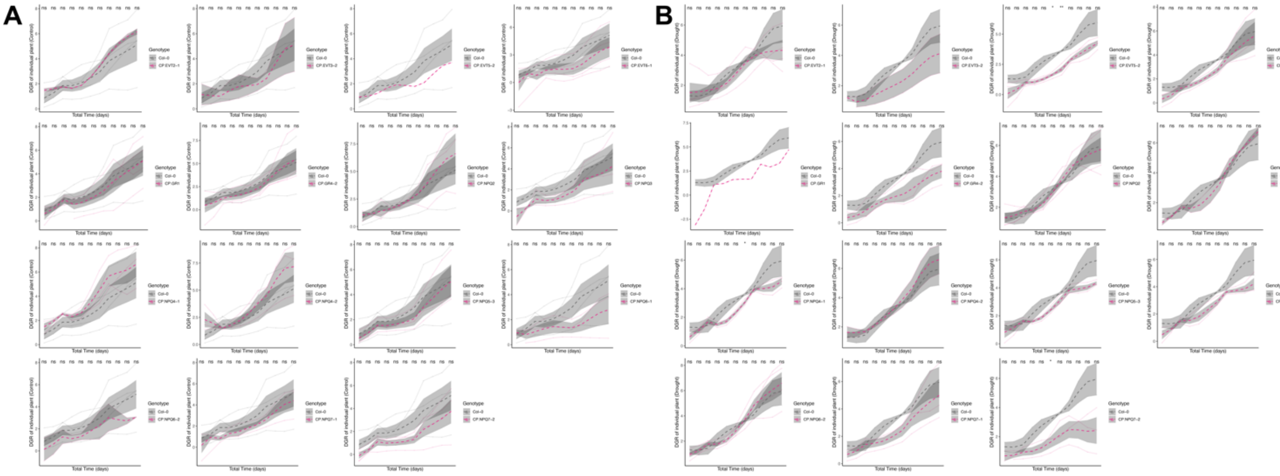

**Figure S7. Second experimental batch of drought-induced changes in daily growth rate among Arabidopsis T-DNA insertion lines.** The Arabidopsis T-DNA insertion lines (*Table S9*) were germinated on agar plates alongside Col-0, and transplanted to soil 1 week after germination. Every 2nd day the pots were weighted and watered to target weight corresponding to 60 and 10% of soil water-holding capacity for control and drought stress treatment respectively. The daily growth rate (DGR) was calculated over 16 h for each individual day of the experiment, and compared to Col-0 for plants grown under **(A)** Control or **(B)** drought stress conditions. The transparent lines represent the data recorded of individual replicates, while the genotype-mean and standard error are represented by the dashed line and grey ribbon respectively. The significant differences between Col-0 and each T-DNA insertion line were evaluated using a t-test, and *, **, ***, and **** signify p-values below 0.05, 0.01, 0.001, and 0.0001 respectively.

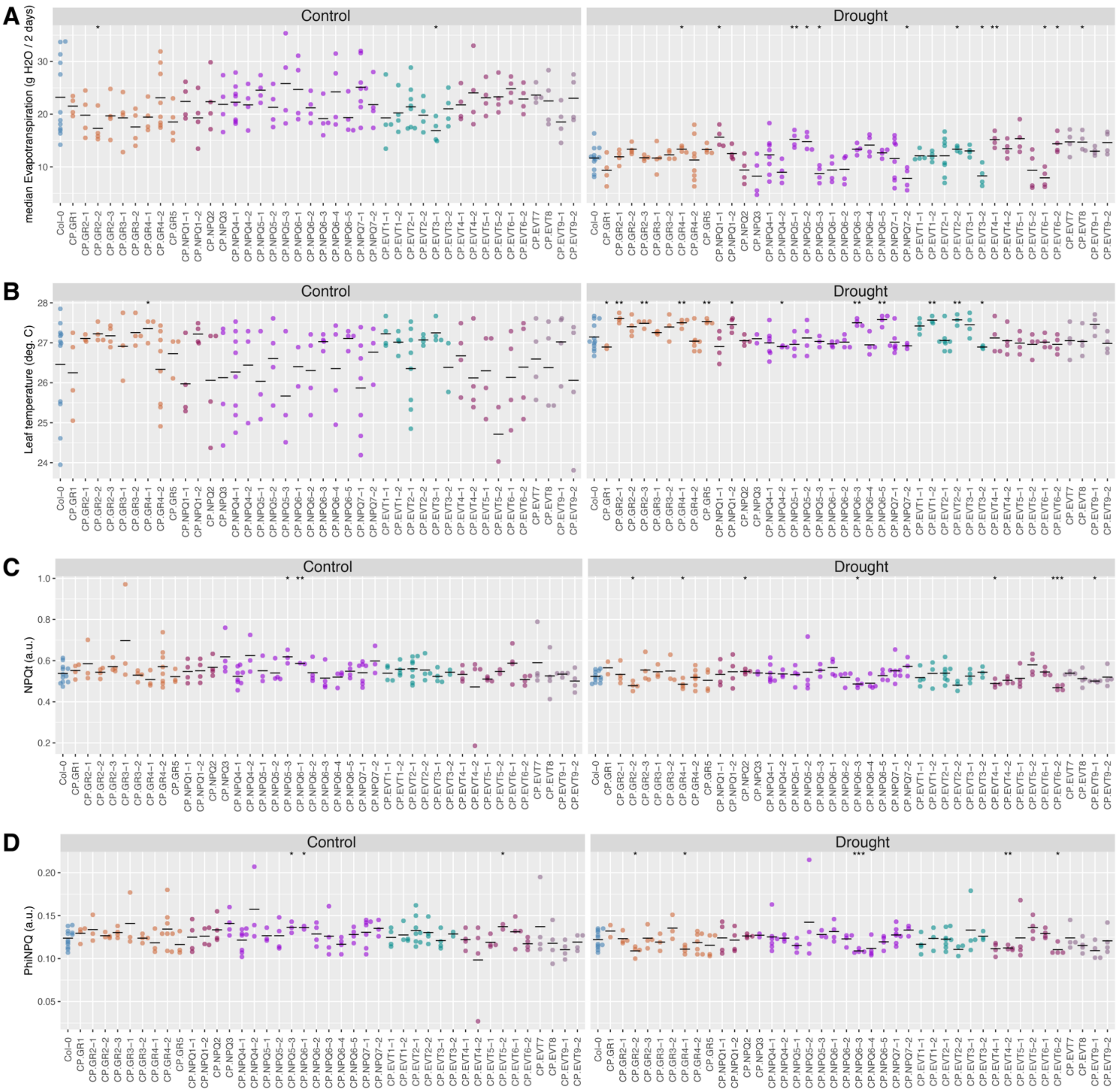


**Figure S8. Drought-induced changes in evapotranspiration, leaf temperature and non-photochemical quenching among Arabidopsis T-DNA insertion lines.** The Arabidopsis T-DNA insertion lines (*Table S9*) were germinated on agar plates alongside Col-0, and transplanted to soil 1 week after germination. Every 2nd day the pots were weighed and watered to target weight corresponding to 60 and 10% of soil water-holding capacity for control and drought stress treatments respectively. **(A)** The median evapotranspiration was calculated per plant over the course of the entire experiment (2 weeks), while **(B)** leaf temperature and **(C-D)** non-photochemical quenching (NPQ) were evaluated at the last day of the experiment (2 weeks after treatment imposition). The individual points represent the data recorded of individual replicates, while the genotype-mean is represented by the horizontal line. The significant differences between Col-0 and each T-DNA insertion line were evaluated using t-test, and *, **, ***, and **** signify p-values below 0.05, 0.01, 0.001, and 0.0001 respectively.

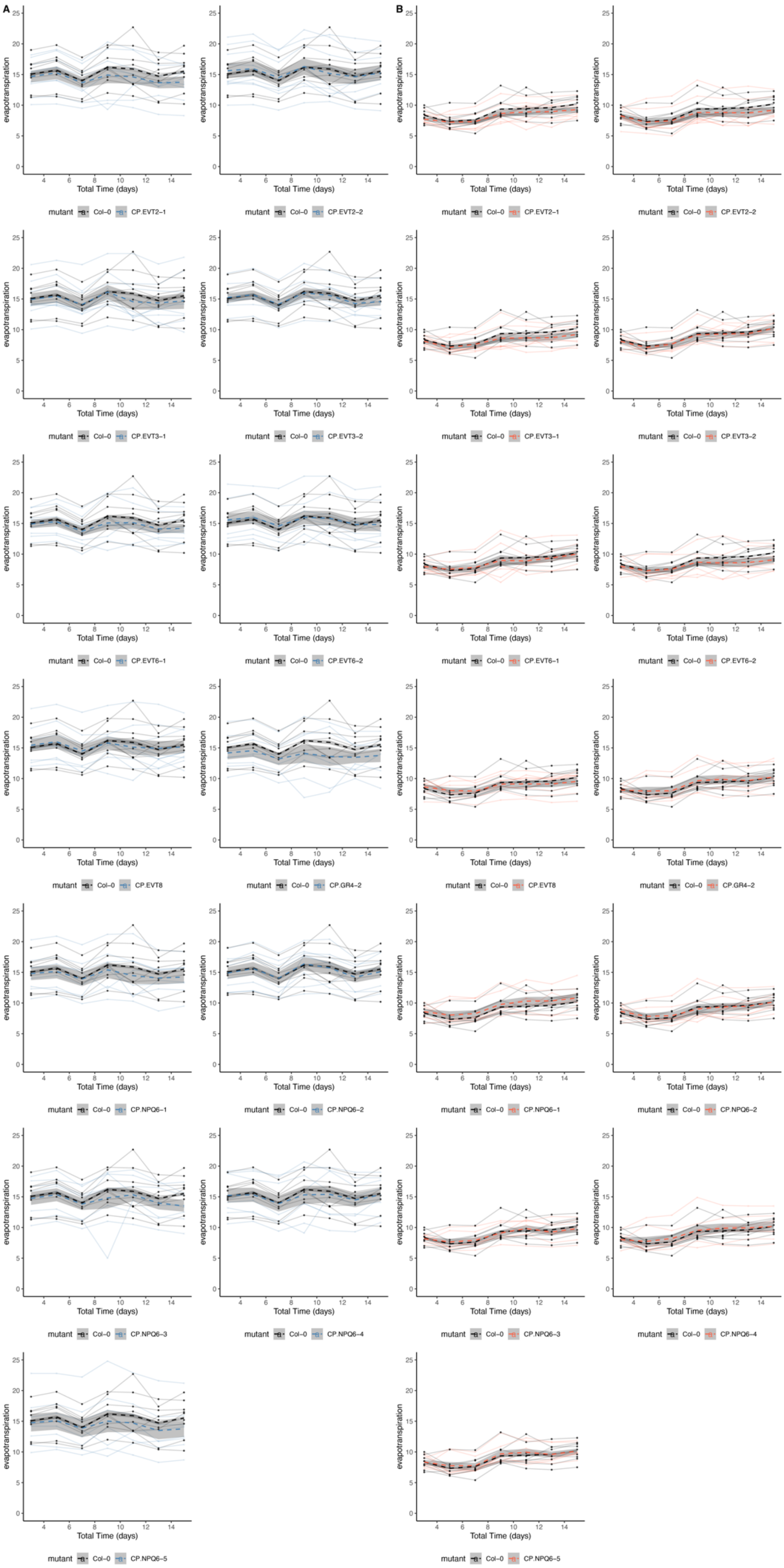
A
B
evapotranspiration
Total Time (days)
mutant Col–0 CP.EVT2–1
mutant Col–0 CP.EVT2–2
mutant Col–0 CP.EVT3–1
mutant Col–0 CP.EVT3–2
mutant Col–0 CP.EVT6–1
mutant Col–0 CP.EVT6–2
mutant Col–0 CP.EVT8
mutant Col–0 CP.GR4–2
mutant Col–0 CP.NPQ6–1
mutant Col–0 CP.NPQ6–2
mutant Col–0 CP.NPQ6–3
mutant Col–0 CP.NPQ6–4
mutant Col–0 CP.NPQ6–5

**Figure S9. Evapotranspiration of mutants versus Col-0.** Evapotranspiration was tracked by recording the amount of water needed to reach target weights every second day for two weeks. (**A)** compares each mutant to Col-0 while under control conditions (60% soil water-holding capacity), whereas **(B)** compares each mutant to Col-0 while under drought conditions (20% soil water-holding capacity). A t-test was performed to identify significant differences between Col-0 and each mutant. No significant differences in evapotranspiration were found between the mutants and Col-0 under both control and drought at any of the studied time points. The color coding for genotypes is reported below each graph. Each transparent line represents the evapotranspiration of one individual plant, and dashed lines and shaded areas represent the genotype average and standard deviation over time respectively.

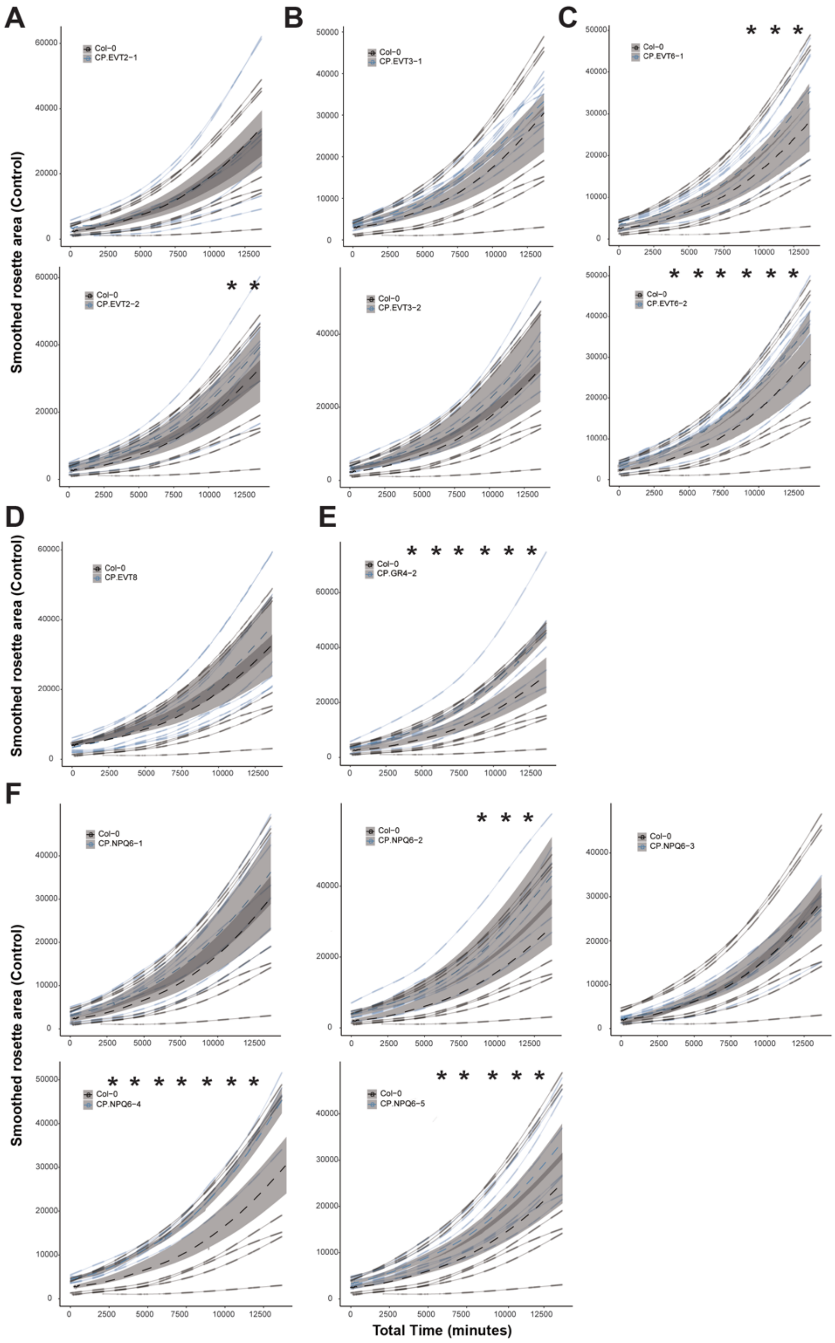


**Figure S10. The growth rate of studied Arabidopsis mutant lines under non-stress conditions.** The mutants target **(A)** 1,8-cineole synthase (EVT2) **(B)** alpha carbonic anhydrase 7 (EVT3) **(C)** WRKY70 (EVT6) **(D)** CAAX amino terminal protease family protein (EVT8) **(E)** xyloglucan endotransglucosylase/hydrolase 16 (GR4) and **(F)** Pentatricopeptide repeat (PPR) superfamily protein (NPQ6). The list of specific mutants can be found in **Supplemental Table S9.** Growth of Col-0 and individual mutant lines recorded under control conditions (i.e. 60% soil water-holding capacity). Transparent lines represent the growth of individual lines, whereas dashed lines represent the genotype average. The shaded surfaces represent the standard error. Significant differences between wild-type and mutant are indicated with *, **, ***, and **** for t-test p-values < 0.05, 0.01, 0.001, and 0.0001 respectively.

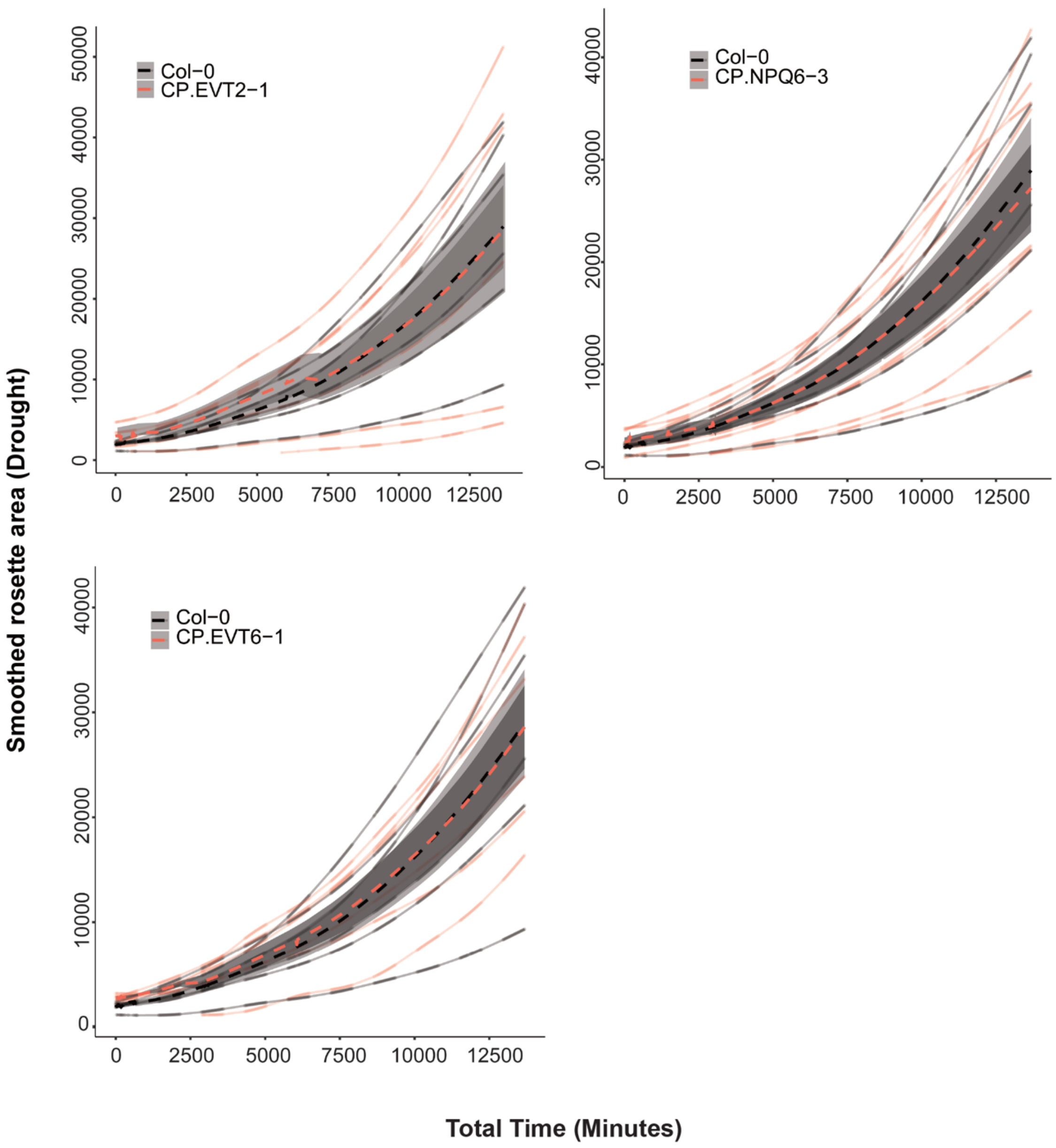


**Figure S11. The growth rate of studied Arabidopsis mutant lines under drought conditions.** The mutant targets including EVT2-1, NPQ6-3, and EVT6-1 were compared to the growth of Col-0 under drought conditions. Transparent lines represent the growth of individual lines, whereas dashed lines represent the genotype average. The pairwise comparison revealed no significant differences between Col-0 relative to each mutant line.
